# Mitochondrial fission is increased in macrophages during mROS production in response to *S. pneumoniae*

**DOI:** 10.1101/722603

**Authors:** Mohammed Mohasin, Katharin Balbirnie-Cumming, Emily Fisk, Elizabeth C. Prestwich, Clark D. Russell, Jennifer Marshall, Clare Pridans, Scott P. Allen, Pamela J. Shaw, Kurt J. De Vos, Christopher J. Hill, Per Bullough, Alison M. Condliffe, Tim J. Mitchell, Helen M. Marriott, David H. Dockrell

**Affiliations:** Department of Infection, Immunity and Cardiovascular Disease, University of Sheffield Medical School, UK; The Florey Institute for Host-Pathogen Interactions, University of Sheffield Medical School, UK; Department of Infection Medicine and MRC Centre for Inflammation Research, University of Edinburgh, UK; Department of Neuroscience, Sheffield Institute for Translational Neuroscience, University of Sheffield Medical School, UK; Centre for Membrane Interactions and Dynamics, Department of Molecular Biology and Biotechnology, University of Sheffield, UK; Department of Biomedical Science, Department of Molecular Biology and Biotechnology, University of Sheffield, UK; Krebs Institute, Department of Molecular Biology and Biotechnology, University of Sheffield, UK; Institute of Microbiology and Infection, School of Immunity and Infection, University of Birmingham, UK

## Abstract

Immunometabolism and regulation of mitochondrial reactive oxygen species (mROS) are critical determinants of the immune effector phenotype of differentiated macrophages. Mitochondrial function requires dynamic fission and fusion, but whether effector function is associated with altered dynamics during bacterial responses is unknown. We show that macrophage mitochondria undergo fission after 12 h of progressive ingestion of live *Streptococcus pneumoniae* (pneumococci). Fission is associated with progressive reduction in oxidative phosphorylation but increased mROS generation. Fission is enhanced by mROS production, PI3Kγ signaling and by cathepsin B, but not by inflammasome activation or IL-1β generation. Reduced fission following PI3Kγ or cathepsin B inhibition is associated with reduced mROS generation and bacterial killing. Fission is associated with Parkin recruitment to mitochondria, but not mitophagy. Fission occurs upstream of apoptosis induction and independently of caspase activation. During macrophage innate responses to live bacteria mitochondria shift from oxidative phosphorylation and ATP generation to mROS production and microbicidal responses with a progressive shift towards mitochondrial fission.

## INTRODUCTION

Macrophages are an essential component of innate immune responses to pathogenic bacteria (1). Pathogen clearance by macrophages involves a co-ordinated response requiring phagocytosis of bacteria, generation of microbicidals and orchestration of the inflammatory response via cytokine/chemokine networks. Macrophage activation results from activation of diverse pattern recognition receptors by microbial products (2, 3). Our understanding of the exact microbicidal strategies utilized by macrophages against extracellular bacteria is still incomplete but we have recently shown that mitochondrial reactive oxygen species (mROS) are an integral part of the anti-bacterial response required to clear *Streptococcus pneumoniae* (pneumococci) (4). *S. pneumoniae* remains the leading cause of bacterial community-acquired pneumonia (CAP) and is a significant cause of bacteraemia and meningitis (2). Host-pathogen interactions between pneumococci and differentiated macrophages provides an informative model for the interaction between extracellular encapsulated bacteria and tissue macrophages (1).

Mitochondria play key roles in innate host defence and contribute to pathogen sensing, cytokine production and pathogen clearance (5, 6). In particular, mitochondria have emerged as critical effectors of delayed-phase microbicidal responses to ingested bacteria, through generation of mROS (4, 7, 8). The emerging area of immunometabolism links metabolic pathway utilization to effector phenotype and has been particularly informative in defining mechanisms underpinning macrophage responses to microbial stimuli (9). Mitochondrial function is regulated by dynamics with a balance between fission and fusion essential for optimal function (10). To date there has been little exploration of how mitochondrial dynamics are altered during microbicidal responses and increased production of mROS. A shift to glycolytic metabolism is associated with mitochondrial fission (11, 12). This is noteworthy since the shift to glycolytic metabolism is recognised as a critical determinant in shaping macrophage activation and innate immune effector function in response to pattern recognition receptor (PRR) stimulation by pathogen-associated molecular patterns (PAMPs) such as LPS (13–15). There is, however, limited understanding of immunometabolic responses and changes in mitochondrial dynamics during challenge with live bacteria, as opposed to microbial components, and how these relate to mROS generation and bacterial killing.

We demonstrate that the host response to pneumococci in macrophages involves induction of fission in response to live bacteria. Fission is associated with reduced oxidative phosphorylation but enhanced mROS production, which promotes fission. Inhibition of phosphoinositide 3-kinase (PI3K) signalling and cathepsin B reduce mitochondrial fission and intracellular bacterial killing. Fission leads to Parkin activation but is not associated with mitophagy and occurs upstream of apoptosis induction. Overall this suggests mitochondrial fission is contemporaneous with mitochondrial adaption to microbicidal function following ingestion of bacteria. (This article was submitted to an online preprint archive (16)).

## RESULTS

### Sustained bacterial exposure results in increased mitochondrial fission in macrophages

We have previously shown that differentiated macrophages have an extensive mitochondrial volume in comparison to less differentiated monocytes and macrophages (17). As shown in Fig. 1A-C, the mitochondrial network showed extensive branch-points in mock-infected BMDM but, in contrast, showed limited complexity after extended bacterial challenge, using a previously developed algorithm for the calculation of branch points (18) as illustrated in Fig. S1. Analysis of the kinetics of this response showed increased mitochondrial fission was a response that progressed over time and was significant by 12-14 h following bacterial challenge (Fig. 1D). When we looked at mitochondrial ultrastructure in BMDM we also noted reduction in cristae following bacterial challenge (Fig. 1E-F).

**Fig 1.**
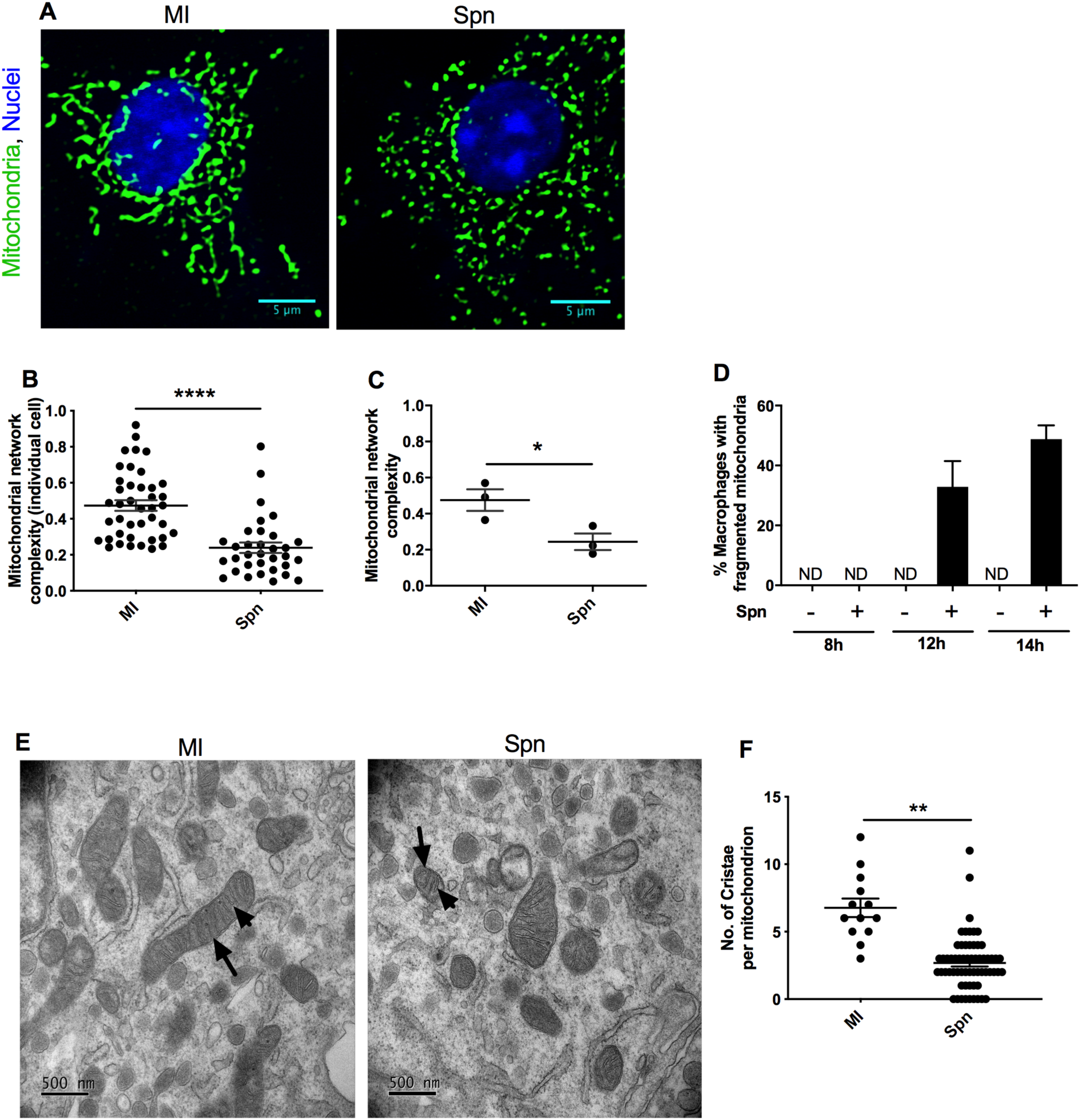
Sustained exposure to *S. pneumoniae* results in increased mitochondrial fragmentation. Bone marrow derived macrophages (BMDMs) were mock-infected (MI) or exposed to *S. pneumoniae* (Spn) at a MOI of 10, for 8-14 h. (A-D) Confocal microscopy was performed after staining with the mitochondrial outer membrane specific marker TOMM20 (green) to delineate mitochondrial structure. (A) Representative filtered images of mitochondrial structure in BMDMs 12 h after exposure under each experimental condition, scale bars = 5 µm, from one of three independent experiments; (B) calculated mitochondrial network complexity (n≥30 cells per condition) across the three independent experiments at 12 h; (C) average calculated mitochondrial network complexity under each condition in the three independent experiments at 12 h; (D) percentage of macrophages with fragmented mitochondria at 8, 12 and 14 h post-bacterial challenge (Spn+) or MI (Spn-), calculated from three independent experiments. (E-F) MI or Spn BMDMs 12 h after challenge were also imaged by transmission electron microscopy. (E) Representative electron micrographs, from one of three independent experiments are shown, scale bars = 500nm. Arrows show mitochondria and arrowheads show cristae; (F) number of cristae per mitochondrion in representative slices were calculated (MI, n=13, Spn, n=62). Data are shown as mean ± SEM. Statistical analysis was performed with one-way ANOVA and Bonferroni post-hoc test (B-C), student paired *t* test (F). *p<0.05, ** p<0.01, ****p<0.0001. ND=not detected.

### Mitochondrial fission is a response to live bacteria

Prior reports have described Drp-1 independent mitochondrial fission in HeLa cells containing *Listeria monocytogenes* (19). In this model fission was induced by the cholesterol-dependent cytolysin (CDC), listeriolysin O, as well as by related CDCs, including pneumolysin expressed by pneumococci, but occurred rapidly after exposure and was transient in duration (20). We addressed the microbiologic requirements for the delayed fission we observed in macrophages. Live bacteria induced fission, but heat-killed bacteria did not (Fig. 2). A pneumolysin deficient mutant at a comparable MOI to the wild-type strain induced lower levels of mitochondrial fission, but by increasing the MOI of the pneumolysin deficient mutant we were able to reconstitute mitochondrial fission to comparable levels, despite absence of the toxin. Exogenous pneumolysin also induced mitochondrial fission only in association with higher lytic concentrations, as confirmed by red blood cell lysis assay (21).

**Fig 2.**
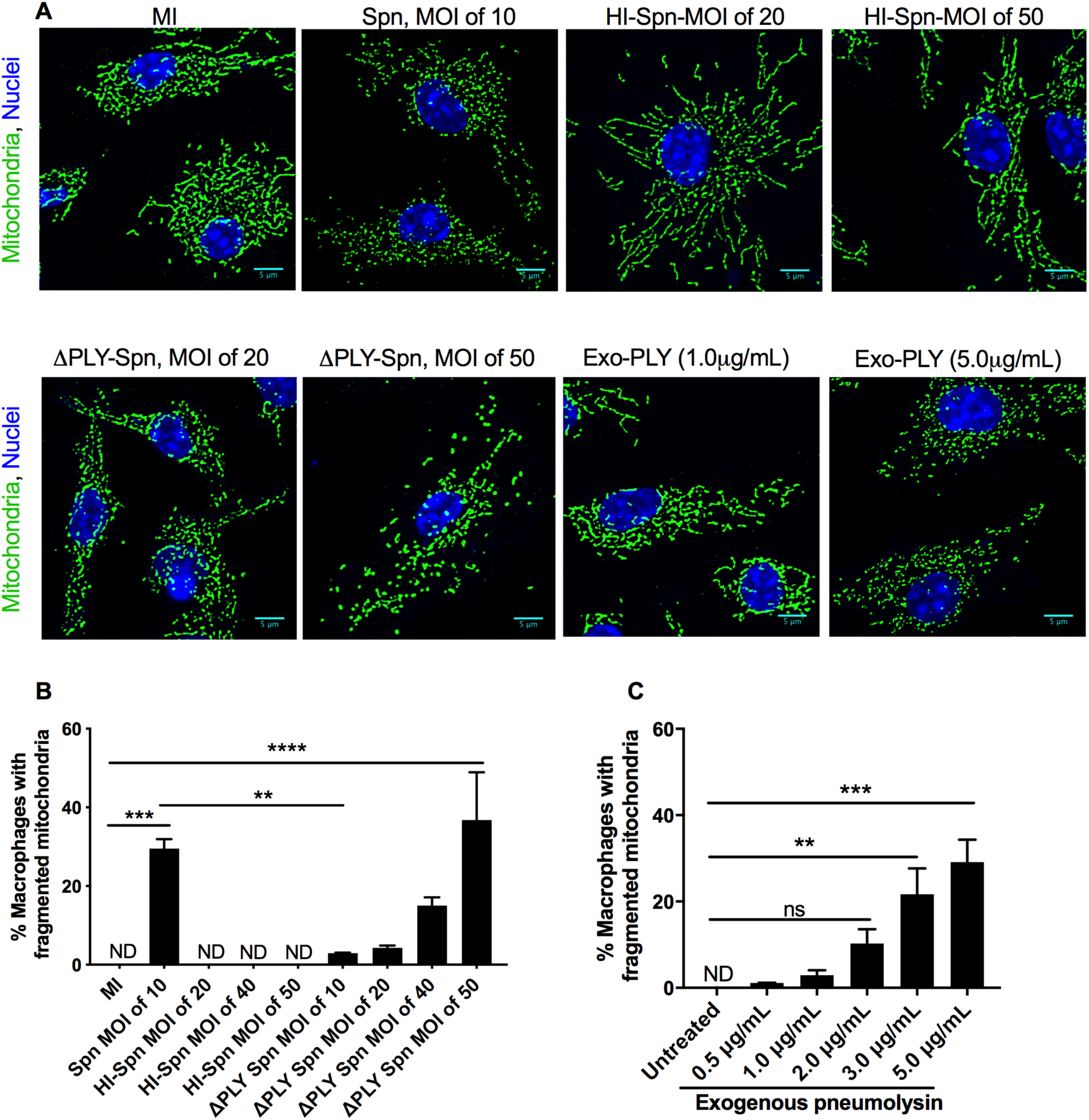
The pore-forming toxin pneumolysin is not essential to trigger increased mitochondrial fission. BMDMs were mock-infected (MI), challenged with *S. pneumoniae* (Spn), heat inactivated Spn (HI-Spn) or a pneumolysin deficient Spn mutant (ΔPLY Spn), at the indicated multiplicity of infection, or alternatively with exogenous pneumolysin (0.5 µg/mL-5 µg/mL) for 12 h. Confocal microscopy was performed after staining with the mitochondrial outer membrane specific marker TOMM20 (green) to delineate mitochondrial structure or with Draq5 to label nuclei (blue). (A) Representative filtered images of mitochondrial structure in BMDMs under a range of conditions, scale bars = 5 µm, from one of three independent experiments. Percentage of macrophages with fragmented mitochondria after challenge with (B) each strain of bacteria or (C) each concentration of pneumolysin. Data are shown as mean ± SEM, (n=3). Statistical analysis was performed with one-way ANOVA and Sidak’s multiple comparison (B) or Bonferroni post-hoc test (C), * p<0.05, ** p<0.01, ****p<0.0001. ND=not detected.

Collectively this suggested that although pneumolysin contributes to mitochondrial fission it is not essential. This therefore emphasised some important features of fission in macrophages responding to live bacteria as opposed to prior reports in other cell types (20). In particular it suggested that in macrophages mitochondrial fission was a response to a bacterial pathogen we have previously shown is contained within the phagolysosome (4), and did not have an absolute requirement for expression of a CDC. This raised the possibility that it arose as a host response to live bacteria.

### Mitochondrial fission is associated with altered mitochondrial metabolism

We next investigated whether fission induced by challenge with live bacteria was associated with changes in cell metabolism, since prior reports suggest reduction of oxidative phosphorylation is associated with enhanced mitochondrial fission (22). Extracellular acidification was enhanced following bacterial exposure (Fig. S2A-B). Basal oxygen consumption (OCR) rate was not significantly altered by bacteria (Fig. S2C) but maximal and ATP linked OCR were reduced after pneumococcal challenge (Fig. 3A-D), as was respiration reserve (Fig. S2D). Moreover, proton leakage (Fig. 3E) and non-mitochondrial OCR (Fig. S2E) were increased by pneumococcal challenge. These findings were also confirmed in human monocyte-derived macrophages (MDM), with confirmation of increased ECAR early after bacterial challenge (Fig. S3A). In addition, a reduction in maximal and ATP-linked (but not basal) OCR and an increase in proton leakage was apparent after bacterial challenge (Fig. S3B-G). Although the shift to glycolytic metabolism was apparent by 4 h the changes in maximal OCR and proton leak were only apparent at the later 16 h time point after bacterial challenge (Fig. S3B-G). Proton leak is often a step designed to limit mROS production when mROS generation is high (23) and in association with this we observed increased mROS generation (Fig. 3F-G). The reductions in maximal and ATP-linked OCR after bacterial challenge were reduced in the presence of mitoTEMPO, an inhibitor of mROS (24), whereas the increases in glycolytic metabolism and non-mitochondrial OCR were not altered (Fig. 3B-D and Fig. S2). Since enhanced mROS generation was a prominent response and contributed to the reduction in changes in OCR we also tested to what extent fission was a direct result of mROS production and documented that mROS inhibition reduced mitochondrial fission after bacterial challenge (Fig. 3H-I). Collectively these results showed that enhanced mitochondrial fission following pneumococcal challenge is associated with a decline in oxidative phosphorylation and an increase in mROS generation. Of note, mROS directly contributes to mitochondrial fission and the alteration in oxidative phosphorylation, while the delayed emergence of the altered parameters of oxidative phosphorylation matches the delayed kinetics of mitochondrial fission.

**Fig 3.**
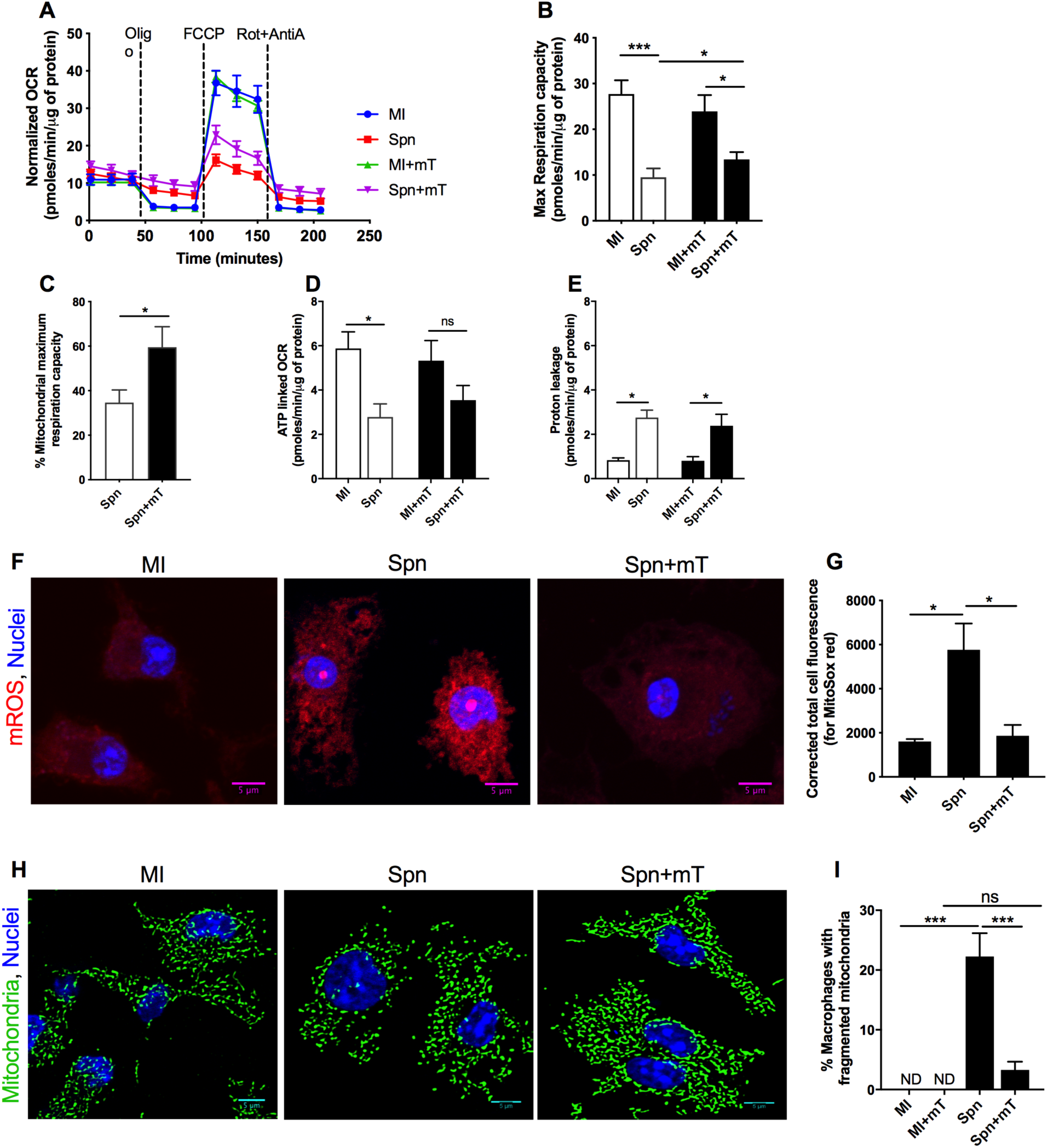
mROS contributes to increased mitochondrial fission and metabolic adaption following bacterial challenge. BMDMs were mock-infected (MI) or challenged with *S. pneumoniae* (Spn) for 12 h after pre-treatment with the mROS inhibitor MitoTempo (+mT) or vehicle control. Subsequently, the mitochondrial oxygen consumption rate (OCR) was measured by the Seahorse X24 extracellular flux analyser. The ATP synthase inhibitor oligomycin A (oligo) was added after baseline OCR acquisition, to measure ATP-linked OCR. The maximum respiration capacity was measured by subtracting non-mitochondrial OCR [calculated following treatment with rotenone (Rot) plus antimycin A (AntA)] from Carbonyl cyanide 4-(trifluoromethoxy) phenylhydrazone (FCCP) treated OCR. (A) Representative graph of the OCR kinetic data with each condition from one experiment. From the OCR kinetic data, the mitochondrial maximum respiration capacity (MRC) (B), percentage of mitochondrial MRC after bacterial challenge with or without mT (C), ATP-linked OCR (D) and proton leakage (E) were calculated. (A-E), n=4. (F-G) Under the same conditions BMDMs were stained with MitoSOX Red to detect mROS (red) or nuclei stained with Draq5 (blue) and imaged by confocal microscopy. (F) Representative unprocessed images from one of three independent experiments are shown and (G) corrected total cell fluorescence was calculated, (n=3). (H-I) Under the same conditions mitochondria were stained with anti-TOMM20 (green) and examined by confocal microscopy. (H) Representative filtered images from three independent experiments are shown and (I) the percentage of macrophages with fragmented mitochondria was calculated, (n=300). Data are shown as mean ± SEM. Statistical analysis was performed with one-way ANOVA with Sidak’s post-hoc test for multiple comparisons and student paired *t* test for pair-wise comparison in (C). *p≤0.05, ***p ≤0.001. ND=not detected.

### PI3K regulates early mitochondrial fission and macrophage microbicidal responses

We next addressed factors which regulate fission. Since fission was associated with live bacteria and mROS production we hypothesized that responses linking pathogen sensing to mROS expression in macrophages could contribute. Phosphatidylinositol 3-kinase (PI3K) signalling enhances mROS production in response to microbial factors in macrophages (25) and is activated by a range of microbial stimuli (26). In line with this we observed that while pneumococcal challenge significantly increased mROS generation that in the presence of pan-PI3K inhibitors, mROS levels were lower and were not significantly increased with LY294002 (Pan-PI3Ki) as compared to the pneumococcal challenge with vehicle control at any time point (Fig. 4A-B). We next addressed if a particular PI3K isoform was associated with regulation of fission and observed significant inhibition of mROS with the PI3Kγ isoform, an isoform highly expressed in macrophages (27), albeit to a slightly lower degree than with the pan-PI3K inhibitors (Fig. 4A and C). The pan-PI3K inhibitor completely blocked fission while the PI3Kγ isoform partially blocked fission (Fig. 4D-F). In keeping with microbicidal roles for mROS in delayed microbicidal responses to pneumococci (4), pan-PI3K and PI3Kγ isoform selective inhibitors reduced bacterial killing, with a numerically greater fold increase in viable intracellular bacteria apparent with the pan-PI3K inhibitor (Fig. 4G-H).

**Fig 4.**
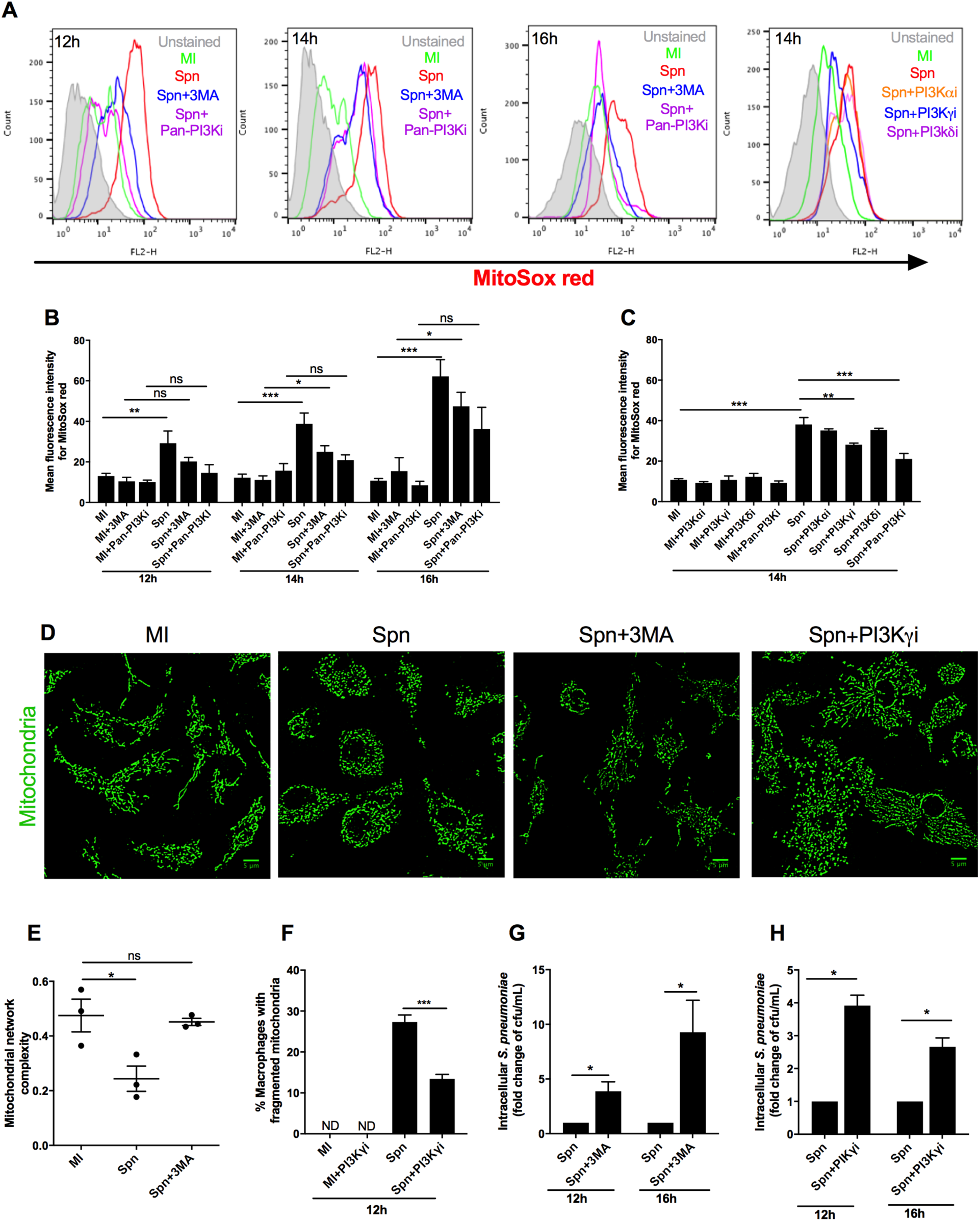
PI3K signaling enhances mROS generation and mitochondrial fission. BMDMs were mock-infected (MI) or challenged with *S. pneumoniae* (Spn) for 12-16 h in the presence of vehicle control or the indicated PI3K inhibitors (3 Methyladenine (3MA), Ly294002 (Pan-PI3Ki), and selective inhibitors of PI3Kα (PI3Kαi), PI3Kγ (PI3Kγi) or PI3Kδ (PI3Kδi) isoforms. (A-C) Cells were stained with MitoSOX Red and analyzed by flow cytometry; (A) shows representative histograms, and mean fluorescence intensity was calculated after PI3K inhibition, n=8 (B) or isoform selective PI3K inhibition, n=3 (C). BMDMs after 12 h of bacterial challenge were stained with anti-TOMM20 to analyze mitochondrial structure. (D) Representative filtered confocal images, scale bars = 5 µm, from one of three independent experiments and mitochondrial network complexity after PI3K inhibition, (E), n=3, and percentage macrophages with fragmented mitochondria after PI3Kγ inhibition (F), n=3 are illustrated. BMDMs were also lysed at selected time points and viable intracellular bacteria (cfu/ml) determined in the presence of 3MA (G), (n=6) or the PI3Kγ inhibitor (H) (n=3) and represented as fold change versus vehicle treatment at each time point. Data are shown as mean±SEM and statistical analysis was performed with one-way ANOVA and Sidak’s post-hoc test and student paired *t* test for pair-wise comparison between two-time points. *p<0.05, **p<0.01, ***p<0.0001, ns- non-significant.

### Cathepsin B regulates early mitochondrial fission and macrophage microbicidal responses

Prolonged exposure to internalized bacteria and mROS leads to lysosomal membrane permeabilization and cathepsin activation (21, 28). Cathepsin B can also stimulate mROS production from mitochondria (29). Lysosomal membrane permeabilization has been linked to mitochondrial fission (30) and we next addressed whether cathepsin B influenced mitochondrial fission. As predicted two separate cathepsin B inhibitors reduced mROS production (Fig.5A-D), but also fission (Fig. 5E-H). In keeping with these findings, both cathepsin B inhibitors also increased viable intracellular bacteria (Fig. 5I-J). The impact of two cathepsin inhibitors on TNFα generation varied from no inhibition to partial inhibition but the effect of cathepsin B inhibitors on generation of IL-1β was more marked with inhibition of greater extent than that observed for TNFα (Fig. S4A-B), in line with previous observations (31). Inhibition of mROS also completely blocked IL-1β production but had no impact on production of TNF-α from macrophages. Since mROS contributes to inflammasome activation resulting in IL-1β generation (28), we also questioned whether inflammasome activation or IL-1β signalling mediated mitochondrial fragmentation and might feedback to mediate the fission associated with mROS or cathepsin B activation. However, the caspase-1 inhibitor YVAD and IL-1RA did not modify mitochondrial fission (Fig. S4C-D).

**Fig 5.**
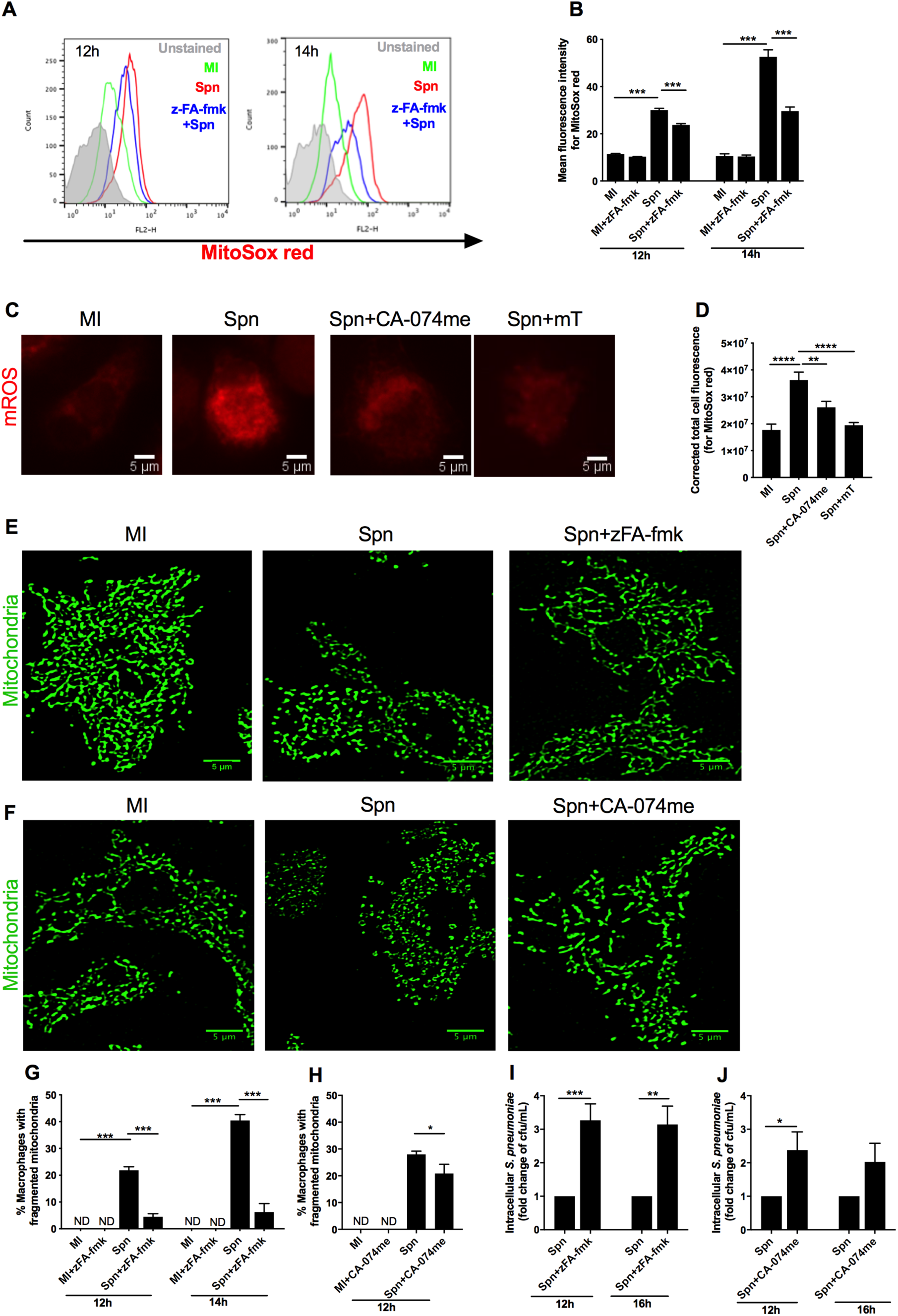
Cathepsin B inhibition modifies mROS generation and mitochondrial fragmentation. BMDMs were mock-infected (MI) or challenged with *S. pneumoniae* (Spn) for 12-14 h in the presence of vehicle control, zFA-fmk or CA-074me. Cells were stained with MitoSOX Red and either analysed by flow cytometry (A-B) or confocal microscopy (C-D). Representative histograms are shown (A) or mean fluorescence intensity plotted (n=4) (B) 12-14h after bacterial challenge in the presence or absence of zFA-fmk treatment. For 14h after bacterial challenge in the presence or absence of CA-074me treatment, representative unprocessed images are shown (C) and (D) corrected total cell fluorescence was calculated, (n=4). BMDMs were stained with anti-TOMM20 (green) and percentage of macrophages with fragmented mitochondria was measured at the indicated time-points 12-14h after bacterial challenge in the presence or absence of zFA-fmk (E, G) or CA-074me (F, H) and representative filtered confocal images, scale bars = 5 µm, from three independent experiments for zFA-fmk (E) and CA-074me (F) at 12 h are shown. Percentage macrophages with fragmented mitochondria 12-14h after bacterial challenge in the presence or absence of zFA-fmk treatment (G), (n=3) or 12h after bacterial challenge in the presence or absence of CA-074me (H), (n=3) are depicted. BMDMs challenged with bacteria for 12-16h in the presence or absence of zFA-fmk were lysed, viable intracellular bacteria (cfu/ml) were calculated and the fold change estimated in the presence of zFA-fmk (as compared to vehicle treatment) (I) (n=6) or were challenged with bacteria for 12-16 h in the presence or absence of CA-074me, cfu/ml calculated and the fold change estimated after CA-074me treatment (as compared to vehicle control) (J) n=3. Data are shown as mean ± SEM. Statistical analysis was performed with one-way ANOVA with Sidak’s multiple comparison test and student paired *t* test for pair-wise comparison **p=0.01, ***p<0.001.

Overall this demonstrated that mROS production and mitochondrial fission were regulated by PI3K signalling and by cathepsin B upstream of mROS roles on inflammasome activation or IL-1β production.

### Mitochondrial fission following bacterial challenge is associated with recruitment of Parkin upstream of apoptosis induction

Having documented that macrophages responding to live pneumococci undergo mitochondrial fission and upregulate mROS production we next examined what the consequences of these processes were to mitochondrial homeostasis. High levels of mROS result in alterations to mitochondrial proteins, which evokes a number of responses including activation of the PINK-1/Parkin system (10). Parkin is an E3 ligase which ubiquitinates damaged mitochondrial proteins to activate their removal via the 26S proteasome system. We found a marked increase in Parkin expression and co-localization of Parkin with mitochondria following bacterial challenge (Fig. 6A-C). In association with this we also observed increased Parkin in the mitochondrial fraction of cells after bacterial challenge (Fig. S5A-C). Since extensive damage to mitochondria can trigger removal by mitophagy (10), we next examined if there was any evidence of mitophagy in macrophages challenged with pneumococci. As shown in Fig. S5D, we found no evidence of activation of the autophagy maker LC3B, although this was induced in macrophages exposed to the mitochondrial oxidative phosphorylation uncoupler Carbonyl cyanide 4-(trifluoromethoxy)phenylhydrazone (FCCP), and also found no evidence for mitophagy associated double-membrane containing vacuoles by transmission electron microscopy (Fig. S5E).

**Fig 6.**
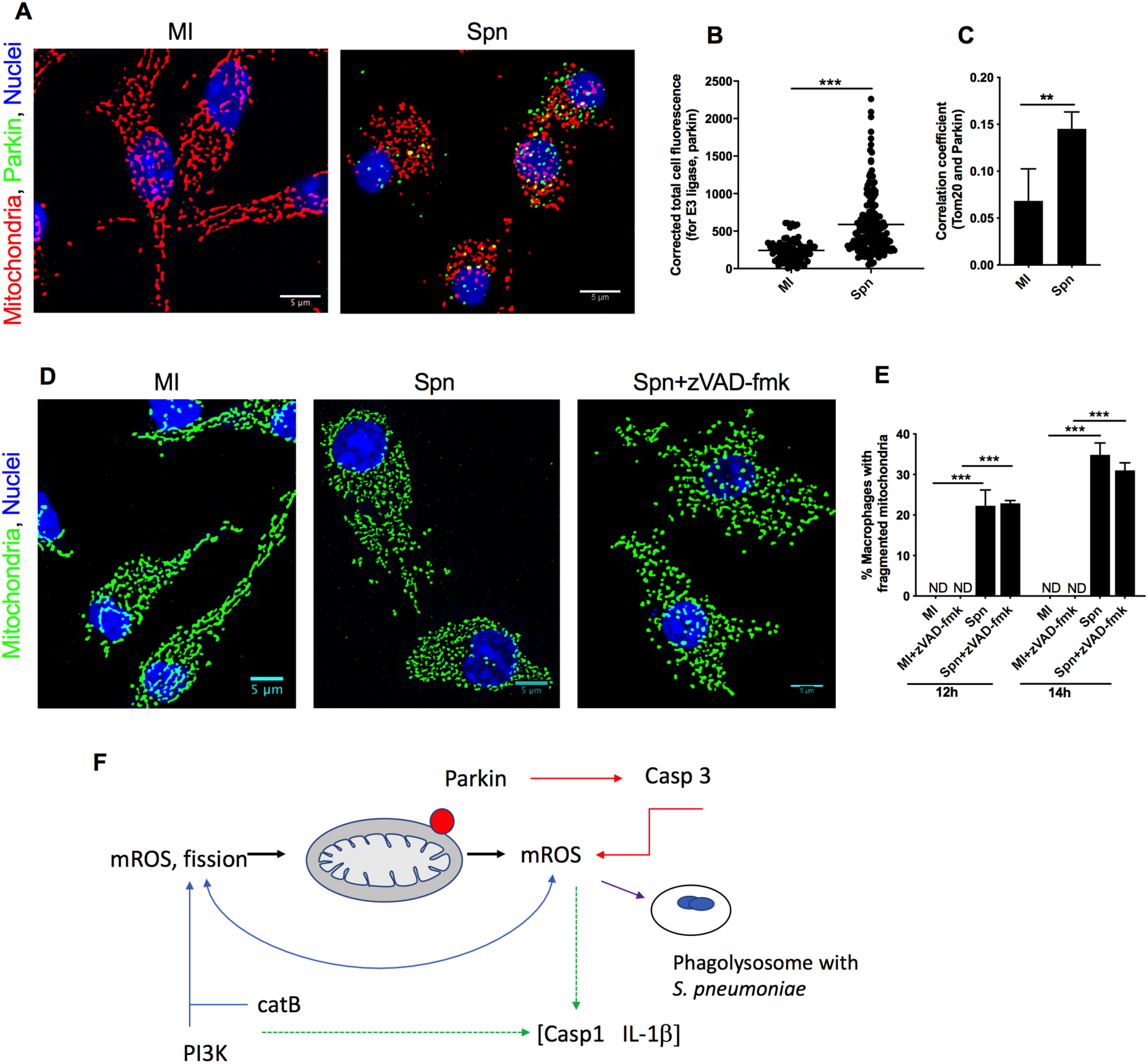
Mitochondrial Parkin recruitment occurs in association with mitochondrial fragmentation before apoptosis. BMDMs were mock-infected (MI) or challenged with *S. pneumoniae* (Spn) for 12 h and stained with anti-TOMM20 (red) to outline mitochondria and with anti-Parkin antibody (green). (A) Representative filtered confocal images, scale bars = 5 µm, from one of three independent experiments are shown. (B) Corrected total cell fluorescence for Parkin was calculated for 78 (MI) or 176 (Spn) BMDMs across three independent experiments and (C) a correlation coefficient was calculated between TOMM20 stained mitochondria and Parkin, (n=3). Data are shown as mean ± SEM. Statistical analyses were performed by paired t-tests. BMDMs (MI or Spn) were also challenged for 12-16 h in the presence or absence of zVAD-fmk and then stained with anti-TOMM20 (green) to outline mitochondrial structure. (D) Representative filtered confocal images of mitochondrial complexity at 12 h, scale bars = 5 µm, from one of three independent experiments are shown and (E) the percentage of macrophages with fragmented mitochondria at 12-14 h are shown, (n=3). Data are shown as mean ± SEM. Statistical analysis was performed with one-way ANOVA with Sidak’s multiple comparison. **p<0.01, ***p<0.001. (F) Schematic Figure illustrating that increased production of mitochondrial ROS (mROS) occurs following increased fission and showing this occurs upstream of subsequent Parkin activation, caspase 3 (Casp3) activation, which further enhances mROS production. Caspase 1 (Casp1) activation and IL-1β production is enhanced by mROS production and cathepsin B (CatB) activation but is downstream of fission and is not related to apoptosis. Phosphoinositide 3-kinase (PI3K) signalling also induces mitochondrial fission and mROS production.

Although a downstream consequence of sustained phagocytosis and bacterial killing is induction of apoptosis, which further increases mROS-dependent killing through caspase-dependent mechanisms (4, 8, 32), we found no evidence of apoptosis when mitochondrial fission was initially observed at the 12 h time point (Fig. S6A-B). Furthermore, a pan-caspase inhibitor zVAD-fmk, which blocks apoptosis-dependent caspases had no impact on fission (Fig. 6D-E), suggesting that mitochondrial fission was upstream of apoptosis induction. Consistent with this, examination of mitochondrial morphology showed that in apoptotic cells mitochondria became swollen and looked similar to those noted after FCCP treatment (compare Fig. S6A with Fig. S5D), while the appearance at the stages prior to apoptosis, at 12-14 h, were characterized by shortening and lack of branching but not by swelling. In addition, loss of inner mitochondrial transmembrane potential (Δψ_m_), a feature of activation of the mitochondrial pathway of apoptosis after pneumococcal challenge (21), was only apparent at relatively low levels at 12 h after bacterial challenge, in contrast to levels at later time points (Fig. S6C-D). This demonstrated that although the recruitment of Parkin was associated with early signs of loss of Δψ_m_ it preceded induction of high levels or advanced stages of apoptosis with caspase activation and occurred early after fission commenced, prior to commitment to apoptosis.

## DISCUSSION

We demonstrate that sustained internalization of live opsonized *S. pneumoniae* results in increased mitochondrial fission in primary macrophages. This is a response that occurs as macrophages reduce reliance on oxidative phosphorylation for ATP generation and as mROS generation progressively increases. mROS, PI3K signalling and cathepsin B promote mitochondrial fission which is associated with increased killing of intracellular bacteria. Ultimately, mitochondrial fission is associated with Parkin activation upstream of apoptosis induction.

Immunometabolism has emerged as a critical determinant of macrophage polarization and phenotype (33). Mitochondrial regulation of macrophage immune function involves release of mROS, which is critical to both pathogen sensing and microbicidal responses (7, 34). Macrophages adapt electron transport chain assembly in response to recognition of live bacteria with associated increases in IL-1β production and fumarate generation, which has been shown to contribute to clearance of *Escherichia coli* and *Salmonella enterica* Typhimurium (35).

Reduced oxidative phosphorylation and increased glycolytic metabolism promote fission (11, 12). Conversely, a putative inhibitor of canonical mitochondrial fission (Mdivi-1) reduced the LPS-induced shift to glycolytic metabolism in murine bone marrow-derived macrophages (BMDM) but as recently highlighted this inhibitor may also directly target complex I and therefore mROS production so does not conclusively establish whether fission is essential for the observed shift in phenotype as opposed to alterations in mROS (36–38). Increased mROS production also enhances fission (10), but fission promotes mROS generation in mitochondria, further amplifying the loop between fission and mROS generation (12). In contrast to the role for mROS in promoting inflammasome activation, fission has been reported to have more variable impact on inflammasome activation. There are reports that it promotes (39, 40) but also inhibits inflammasome activation (41, 42). The shift to glycolytic metabolism during the macrophage host response to bacterial infection would be anticipated to promote mitochondrial fission, reduce reliance on mitochondrial ATP generation and enhance mROS mediated innate immune responses.

Although reports also suggest that microbial factors, in particular cholesterol-dependent cytolysins, may stimulate fission in non-myeloid cells, including pneumolysin produced by *S. pneumoniae* (19, 20), we found that although pneumolysin was a contributory factor, it was not essential for fission. Thus, in macrophages it is live bacteria that promote the delayed mitochondrial fission we observed, similar to the observations of Garaude and colleagues, where transient electron transport chain adaptations involving a shift to complex II activity enable microbicidal responses that also required live bacteria (35). In our case, however, the adaption we observed was a delayed response that progressed over time. We found that PI3K signalling, a known component of pathogen sensing pathways, and in particular the PI3Kγ isoform which is highly expressed in macrophages, promoted fission (27, 43).

PI3Kγ contributes to leukocyte recruitment, activation and nicotinamide adenine dinucleotide phosphate-dependent ROS generation (27, 43), and we suggest an additional role regulating fission and mROS production. PI3K inhibition has also reduced mitochondrial fission in tumor cells, although in this setting this was achieved in association with modulation of trafficking to the cortical cytoskeleton and in association with mROS enhancement (44). We also identified cathepsin B as a factor regulating mitochondrial fission. Cathepsin B is activated and released during phagolysosomal membrane maturation following ingestion of pneumococci (21). It has been shown to stimulate mROS production (29). Both mROS and cathepsin B contribute to inflammasome activation (45, 46), but we found no evidence that caspase 1 activation, or downstream IL-1β production, regulate mitochondrial fission, even though we confirmed cathepsin B contributes to IL-1β release from macrophages. Therefore, we conclude that mROS and cathepsin B induced inflammasome activation occurs downstream of regulation of fission and inflammasome activation, and the resulting IL-1β generation does not form a regulatory feedback loop influencing fission (see Fig. 6F).

A critical factor in fission in our model is mROS or pathways that enhance mROS production. Oxidative stress is known to lead to formation of mitochondrial-derived vesicles and the release of these from mitochondria is dependent on Parkin and PINK-1 (47). It therefore remains to be proven whether the fission we observe which is associated with mROS and Parkin recruitment to mitochondria is regulated by a similar pathway. Ultimately progressive mitochondrial fission and oxidative stress will lead to loss of Δψ_m_ and dysfunctional mitochondria could activate mitophagy (48) or mitochondrial pathways of apoptosis induction via cytochrome c release and mitochondrial outer membrane permeabilization (MOMP) (49, 50). Both processes involve PTEN (phosphatase and tensin homolog) induced putative kinase 1 (PINK1) and Parkin, and the extent of loss of Δψ_m_ and activation of the E3 ligase Parkin determines whether cells undergo mitophagy (51) or apoptosis (52). We observed no evidence of mitophagy, while we have previously documented this model results in apoptosis in association with downregulation of Mcl-1 via ubiquitination and proteasomal degradation (32). We have shown that cathepsin D stimulates Mcl-1 ubiquitination via the E3 ligase MULE (53). Parkin is another potential mediator of Mcl-1 ubiquitination (52). The extent of loss of Δψ_m_ in the face of sustained oxidative stress during macrophage responses to live bacteria is likely to favour apoptosis over mitophagy (21). Fission can occur as an early event in apoptosis (54) and occurs upstream of caspase activation (55). Consistent with this, in our model fission was unaltered by caspase inhibition. However, we found evidence that fission is regulated by specific factors (mROS and cathepsin B), which we have previously demonstrated do not mediate macrophage apoptosis during pneumococcal challenge (4, 53). This suggests that although fission and enhanced mROS play a key role in macrophage microbicidal responses, through intracellular pathogen killing and cytokine generation, they are not required for the execution phase of apoptosis, which proceeds as a result of Mcl-1 downregulation and MOMP despite inhibition of mROS and mitochondrial fission.

In conclusion, we suggest that sustained responses to internalized live bacteria shift mitochondrial dynamics progressively in favour of fission in association with reduced oxidative phosphorylation. This is both the result of mROS production but enables further mROS generation which can enhance IL-1β production and enable microbicidal responses that limit intracellular pathogen survival. Although *S. pneumoniae* is relatively resistant to the direct effects of ROS and the initial effects of mROS may be indirect (56), the subsequent induction of caspase-dependent apoptosis increases mROS generation and enables mROS to act in combination with other microbicidal responses (4), while apoptosis effectively terminates pro-inflammatory cytokine responses. The reliance of macrophages on mROS to effectively control clearance of internalized bacteria requires an effective program that includes mitochondrial fission and adapting mitochondrial function away from oxidative phosphorylation and ATP generation to prioritize microbicidal function.

## MATERIALS AND METHODS

### Ethics Statement

Peripheral blood mononuclear cells were isolated from whole blood donated by healthy volunteers with written informed consent, and differentiated into monocyte derived macrophages (MDMs), as approved by the South Sheffield Regional Ethics Committee (07Q2305) and by the Edinburgh Accredited Medical Regional Ethics Committee (15-HV-013). Animal experiments were performed in accordance with U.K. Government Home Office Regulations (Animals (Scientific Procedures) Act 1986) under Project Licences PPL 40/3726 (Professor David H. Dockrell) and PPL 70/8915 (Dr. Clare Pridans) and ethical approval was granted by the University of Sheffield Local Ethical Review Panel and The University of Edinburgh’s Protocols and Ethics Committees respectively. The animal care and use protocols adhered to National Centre for the Replacement, Refinement and Reduction of Animals in Research guidelines (Responsibility in the use of animals in bioscience research, April 2019 and Animal Research: Reporting of *In Vivo* experiments guidelines, June 2010).

### Bacteria and infection

Serotype 2 *S. pneumoniae* (D39 strain; NCTC 7466) and the pneumolysin deficient strain *S. pneumoniae* (PLYSTOP D39) were cultured and stored as previously described (21, 57). In some experiments bacteria were heat-inactivated by placing in water at 60*°*C for 40 min. Prior to macrophage challenge bacteria were opsonized in RPMI (Sigma-Aldrich) containing either 10% mouse immunized pooled serum or human immunized pooled serum with detectable levels of anti-pneumococcal antibodies for bone marrow-derived macrophages (BMDMs) or human monocyte-derived macrophages (MDMs), respectively. The multiplicity of infection (MOI) or bacteria to macrophage ratio was 10 unless otherwise stated. Endotoxin-free pneumolysin was produced by Prof. Tim Mitchell (University of Birmingham) and shown to be haemolytic in a sheep red blood cell haemolytic assay as previously described, before incubation with macrophages for 12 h at 0.5-5 µg/mL (21). Cultures were incubated for the indicated time before analysis.

### Isolation and culture of macrophages

Mouse BMDM were obtained from C57BL/6 mouse femurs and tibias and differentiated by culture of 0.5 x 10^6^ cells per well in 24 well plates (Corning) on 13 x 13 mm cover slips (VWR International), or alternatively in either T25 or T75 flasks (Corning), in Dulbecco’s Modified Eagle’s Media DMEM;(Bio Whittaker®), Lonza) with 10% fetal calf serum (FCS) with low LPS (HyClone, Thermo Scientific) and 10% conditioned L929 media as a source of CSF-1, as described previously (1). Cells were plated at 2 x 10^6^ cells and cultured in RPMI with 10% FCS with low endotoxin levels (Lonza). After 14 d, macrophages were challenged with bacteria or mock-infected and incubated for 1 h on ice, followed by 3 h at 37*°*C with 5% CO_2_. After washing three times with PBS, cells were incubated for the indicated time period at 37*°*C with 5% CO_2_ before analysis.

In certain experiments, macrophages were incubated with one of the following reagents for 1 h prior to bacterial challenge; 100 µM MitoTempo (Enzo Life) to inhibit mROS, 1 mM 3-methyladenine (3MA) (Sigma), or 15 µM Ly294002 as pan-PI3K inhibitors, 10 µM A66 as a PI3Kα inhibitor, 10 µM A560524 as a PI3Kγ inhibitor, 3 µM IC87114 as a PI3Kδ inhibitor, 50 µM *N*-benzyloxycarbonyl–Phe-Ala fluoromethyl ketone (zFA-fmk) (ApexBio) or 25 µM (L-3-*trans-*(Propylcarbamyl) oxirane-2-carbonyl)-L-isoleucyl-L-proline (CA-074me) (Sigma) to inhibit cathepsin B, 50 µM carbobenzoxy-valyl-alanyl-aspartyl-[O-methyl]-fluoromethyl ketone (zVAD-fmk) (ApexBio) as a pan-caspase inhibitor or 10 µM N-acetyl-tyrosyl-valyl-alanyl-aspartyl chloromethyl ketone (Ac-YVAD-cmk, Calbiochem) as a caspase 1 inhibitor, 200 ng/mL recombinant IL-1RA (Pepro Tech) or 50 ng/mL sTNFR1 (Pepro Tech) prior to challenge with *S. pneumoniae*.

### Confocal immunofluorescence microscopy of mitochondria

To analyze mitochondrial network complexity mitochondrial outer membranes were stained and analyzed by confocal microscopy as previously described (18). After the indicated time periods macrophages were fixed with 4% paraformaldehyde for 20 min at room temperature (RT), permeabilized using 0.1% Triton-X-100 with 50 mM NH_4_Cl for 15 min, and blocked with working solution PGAT (PGAT composition: 0.2% Gelatin with 0.02% Na-azide and 0.01% Triton X-100 in PBS) for 15 min. The blocking solution was replaced with anti-translocase of outer membrane 20 (TOMM20) (rabbit polyclonal IgG (FL-145) or mouse monoclonal IgG2a λ (F10) when staining MDM, (both Santa Cruz) 200 µL/well at a 1:500 dilution in PGAT solution and incubated overnight at 4°C. The following day, cells were washed 3 times with the PGAT solution before being treated with secondary antibody Alexa Fluor 488-conjugated goat anti-rabbit IgG (A11034; Thermo Fisher Scientific), Alexa Fluor 568-conjugated goat anti-rabbit IgG (A11011; Thermo Fisher Scientific) or Alexa Fluor 488 conjugated goat polyclonal anti-mouse IgG (A32723; Molecular Probes) at a 1:500 dilution in PGAT solution and incubated for 1 h at RT. Cells were then washed three times in PGAT and twice in PBS. Nuclei were stained with 5 µM Draq5 (Biostatus Ltd. 1:1000 in PBS). Coverslips were mounted onto microscope slides with a glycerol free poly-(vinyl alcohol) Tris-MWL 4-88 (Citifluor) mounting agent. To assess mitochondrial network complexity, 8-12 Z-stack images per cell from 30 representative cells after TOMM20 staining were acquired with a LSM510 inverted confocal fluorescence microscope (Zeiss) using a 63x1.4 oil objective (zoom 2) lens, 488 nm and 633 nm excitations lasers and 500-530 nm and 660-704 nm emission spectrums for TOMM20 and nuclear stains, respectively. Z-stack images were converted into maximum projected images by ImageJ (v1.8, NIH), as previously described (58). Subsequently, the single cell images were filtered using a 13 x 13 Mexican Hat shaped kernel, a filter defining the edges of the mitochondrial network (59), before being subjected to a Huang threshold to remove background from the image (60). The images were then skeletonized, removing any pixels touching a background pixel, except where removal would result in the breaking of a continuous region of pixels, thus resulting in the formation of a single pixel-wide skeleton. Finally, the binary connectivity was quantified using a binary connectivity plugin (61) (plugins available from http://www.mecourse.com/landinig/software/software.html and adapted by K. J. De Vos (18)) in ImageJ software, as shown schematically in Fig. S1B and C. This plugin generates an output of 0 for background signal, 1 for a single pixel, 2 for an end point signal, 3 for a junction with two neighboring pixels, 4 or more for a branch point with three or more neighboring pixels, as described in the computational model developed by Sukhorukov et al. (62). The network complexity of each individual cell was measured as the ratio of total branch points to total end points. At least 300 representative macrophages were also counted per condition from the acquired images and data recorded as the percentage of macrophages with fragmented mitochondria. Macrophages were scored as having fragmented mitochondria if mitochondria appeared fragmented. Macrophages were observed to show either a regular branched structure or near complete loss of structure with virtually no cells showing partial degrees of fragmentation. Filtered images are shown in Figures in the paper and the corresponding unfiltered images are included in Fig. S7.

For mitochondrial co-staining with E3 ubiquitin ligase Parkin, BMDMs were labeled with rabbit polyclonal anti-TOMM20 (Fl-145) and 1:500 mouse monoclonal anti-Parkin IgG2b κ (PRK8, *#*32282 Santa Cruz) primary antibodies in PGAT solution overnight at 4°C and then Alexa Fluor 568 conjugated goat anti-rabbit and Alexa Fluor 488 conjugated goat anti-mouse secondary antibodies. Nuclei were stained with Draq5 as above. Parkin fluorescence intensity for each condition was measured as the corrected total cell fluorescence (CTCF) by ImageJ as described (4). The Pearson’s correlation coefficient for TOMM20 and Parkin in BMDMs was calculated as described (4) . To allow co-staining with LC3B, human MDM cultures (or positive-controls generated by treatment with 20 µM carbonyl cyanide *p*-triflouromethoxy-phenylhydrazone (FCCP) for 12 h) were labeled with mouse monoclonal anti-TOMM20 (F-10) and 1:500 rabbit polyclonal anti-LC3B (ab48394, Abcam) primary antibodies, and subsequently Alexa Fluor 488 conjugated goat anti-mouse and Alexa Fluor 568 conjugated goat polyclonal anti-rabbit secondary antibodies (63). Nuclei were stained with Draq5.

For mitochondrial ROS (mROS) staining, cells with or without pre-treatment with MitoTempo (Sigma-Aldrich) or CA-074me for 1 h before challenge with bacteria were washed 3 times with pre-warmed Hanks’ Balanced Salt Solution (HBSS, Thermo-Fisher Scientific) and mROS were stained using 2.0 µM MitoSOX Red (Life Technologies) in pre-warmed phenol red free and serum free RPMI media supplemented with 2 mM L-glutamine for 30 min. Cells were fixed with 2% paraformaldehyde (in PBS). Nuclei were stained with Draq5, coverslips mounted onto microscope slides and Z-stack images were acquired, as above, using 488 nm and 633 nm excitations lasers and 565-615 nm (for MitoSOX Red) and 661-704 nm (for Draq5) emission spectrums. For quantification, mROS fluorescence intensity for each condition was measured as the total corrected cell fluorescence, TCCF = integrated density – (area of selected cell - mean fluorescence of control), as previously described (4). Unprocessed images were used for analysis and depicted.

To measure apoptosis 200 µL/well NucView^TM^ 530 red solution (2 µM in phenol red free and serum free RMPI medium) (Biotium) was added for 30 min at 37*°*C and 5% CO_2_. Cells were fixed with 2% paraformaldehyde for 15 min at RT and DAPI (0.5 µg/mL in PBS) was added for 12 min at RT and cells were permeabilized and mitochondria stained as above. The NucView^TM^ 530 (red) positive DAPI (blue) positive cells were counted as apoptotic. At least 300 nuclei were counted for each condition using the Zeiss LSM510 inverted fluorescence microscope.

### Transmission electron microscopy

Mouse BMDMs grown in T25 flasks were challenged with *S. pneumoniae* or mock-infected for 12 h then were washed 3 times with 2 ml Hanks’ Balanced Salt Solution (HBSS) per flask and accutase was added for 15 min at 37°C and 5% CO_2_. The cell suspension was centrifuged at 2000g for 10 min, washed once with HBSS and centrifuged again at 2000g for 10 min. The pellet was fixed overnight with 2.5% glutaraldehyde in 0.1 M sodium phosphate buffer at 4°C. The pellet was then washed twice with 0.1 M sodium cacodylate buffer for 30 min at 4°C. Secondary fixation was carried out in 2% aqueous osmium tetroxide for 2 h at room temperature (RT) as previously described (68), then washed free from secondary fixative twice with 0.1 M sodium cacodylate buffer for 30 min before being dehydrated through a graded series of ethanol solutions in water; 75%, 95%, and 100%, and 100% ethanol dried over anhydrous copper sulphate for 15 min each at RT. The pellet was cleared of ethanol in an intermediate solvent of propylene oxide for 15 min, twice, at RT, before being infiltrated by in a 50/50 mixture of propylene oxide/araldite resin overnight at RT on a rotor mixer in a fume hood. The following day this mixture was mostly discarded, the remainder propylene oxide evaporated in the fume hood leaving the pellet which was then placed in full strength araldite resin with one change, for 6-8 h at RT.

Subsequently the specimen pellet was embedded in fresh araldite resin and cured at 60°C for 48-72 hrs. Ultrathin (∼85 nm) sections were cut on a Reichert Ultracut E ultramicrotome with a diamond knife, transferred to copper grids and stained with uranyl acetate and lead citrate. Electron micrographs were recorded at 80 kV at a nominal magnification of 13,000x on a FEI Tecnai G2 Biotwin Spirit microscope equipped with a Gatan Orius digital camera. The mean number of cristae per mitochondria for 13-62 mitochondria was recorded for each condition.

### Metabolic measurements

Macrophages grown in T75 flasks for 14 d were washed once with sterile Dulbecco’s PBS (Life Technologies) and treated with accutase (Biolegend) for 15 min at 37*°*C and 5% CO_2_. The detached cells were re-seeded in XF24 cell plates (Agilent Technologies) at 150,000 MDMs/well or 200,000 BMDMs/well. Cultures were then challenged with bacteria or mock-infected in the presence or absence of MitoTempo and at the indicated time points cultures were washed twice with XF medium supplemented with 4.5 g/L D-glucose, 2.0 mM L-glutamine, 1.0 mM sodium-pyruvate, 100 U/L penicillin and 100 µg/mL streptomycin at pH 7.4 (adjusted with 1.0 M NaOH). Next 630 µL modified XF medium was added to each well and incubated for 1 h at 37^°^C without CO_2_. At the same time the XF24 utility plate, containing sensors probes, was set up using a plate previously submerged and incubated in XF calibrant (Agilent Technologies) overnight at 37*°*C. 70 µL oligomycin A (15 µM), 77 µL FCCP (20 µM) and 85 µL rotenone (10 µM) plus antimycin A (10 µM) (all Sigma-Aldrich) were added to the cartridge injection ports A, B and C, respectively and incubated for 1 h at 37*°*C without CO_2_. Finally, the calibration plate was loaded into the XF24 flux analyser. After calibration completion, the utility plate without cartridge was unloaded and the cell plate was loaded into the XF24 analyser (Seahorse, Agilent Technologies). After equilibration, the cartridge containing the oxygen sensor, measuring the oxygen consumption rate (OCR) and the cartridge containing the proton sensor, measuring the extracellular acidification rate (ECAR) kinetics were run before and after injecting oligomycin, FCCP and rotenone plus antimycin A, respectively. The key parameters of mitochondrial oxidative phosphorylation and cytosolic glycolysis were calculated from OCR and ECAR, respectively, as described in (64). The ATP synthase inhibitor oligomycin A (oligo) was added after baseline OCR acquisition, to measure ATP-linked OCR. The maximum respiration capacity was measured by subtracting non-mitochondrial OCR [calculated following treatment with rotenone (Rot) plus antimycin A (AntA)] from Carbonyl cyanide 4-(trifluoromethoxy) phenylhydrazone (FCCP) treated OCR. Data were normalized by total protein.

### Flow cytometry

Mitochondrial specific ROS was also measured by flow cytometry. Macrophages were stained using 2.0 µM MitoSOX Red (Life Technologies) diluted in pre-warmed phenol red free RPMI media supplemented with 2mM L-glutamine and incubated for 30 min at 37*°*C and 5% CO_2_. As a positive control, BMDM were treated with rotenone (2.0 µM) and antimycin A (10 µM) for 30 min at 37^°^C and 5% CO_2._ Loss of mitochondrial inner transmembrane potential (Δψ_m_) was measured with 10 µM of 5,5′,6,6′-tetrachloro-1,1′,3,3′-tetraethyl benzimidazolyl carbocyanine iodide (JC-1; eBioscience), diluted in phenol red free RPMI media supplemented with 2 mM L-glutamine) (eBioscience, 65-0851-38) for 30 min at 37*°*C with 5% CO_2._ Cells were re-suspended in PBS (300 µL/well), after gentle scrapping and washing 3 times with HBSS. Both oxidized MitoSOX Red and the loss of JC-1 aggregates as a marker of loss of Δψ_m_ were measured using a FACS Calibur (Becton Dickinson) via the FL-2H channel. The forward and side scatters were used to distinguish cell populations and a total of 10,000 events were recorded. The instrument settings were saved and used for subsequent sample acquisition and analyses. Data were analyzed using FlowJo software, version 8.8.4 (Tree Star Inc.).

### Cytokine assay

IL-1β and TNF-α levels in the culture supernatants were measured using mouse IL-1β and TNF-α DuoSet enzyme-linked immunosorbent assay (ELISA) kits, respectively (eBioscience, Hatfield, United Kingdom) according to the manufacturer’s specifications.

### Intracellular killing assay

The intracellular killing assays were carried out as previously described (8). To perform modified gentamicin protection assays cells were treated at the indicated time points with 20 µg/mL gentamicin and 40 U/mL penicillin G (Sigma, PENNA-100MU) for 30 min at 37*°*C and 5% CO_2_, to kill extracellular bacteria. Subsequently cells were washed 3 times with PBS before being treated with 2% saponin (250 µL/well) (Sigma) in distilled water and incubated for 15 min at 37*°*C and 5% CO2. In some experiments 12 h and 16 h killing assays, macrophages were performed in the presence or absence of the indicated inhibitors or vehicle controls. In these experiments the appropriate reagent was added 1 h before bacterial challenge, added again at the time of bacterial challenge and at the 4 h time point after bacteria were washed off. For 16h killing assays, macrophages were ‘pulsed’ with gentamicin and penicillin G at 12 h as above, then ‘chased’ with 0.75µg/mL vancomycin (Sigma) in the presence of the appropriate reagent for a further 4 h before washing and saponin lysis (8).To perform surface viable counts, 750 µL PBS was added to each well and macrophages were lysed with vigorous pipetting and scraping across the wells. Subsequently the lysates were serially diluted and plated on Columbia blood agar plates. Data were calculated as cfu/mL and ratios of inhibitors compared to vehicle controls.

### SDS-PAGE and Western blotting

Macrophages cultured in T25 flasks were challenged with bacteria or mock-infected and cytosolic or mitochondrial fractions isolated as described previously (65). Protein was quantified using a modified Lowry protocol (*DC* protein assay; Bio-Rad Laboratories), and equal protein was loaded per lane, 20 µg of protein from cytosolic and 10 µg from mitochondrial fractions. Samples were separated by SDS-PAGE (12%) and blotted onto poly-vinylidene difluoride (PVDF) membranes (Bio-Rad Laboratories) with protein transfer confirmed by Ponceau S staining. Blots were incubated overnight at 4°C with antibodies against Parkin (PARK8, mouse monoclonal, Santa-Cruz Biotechnology Inc., cat no. SC-32282, 1:200 dilution in 5% milk TBS-tween), Actin (Rabbit polyclonal, Sigma-Aldrich 1:10000 dilution) or Voltage dependent anion channel (VDAC) (Rabbit polyclonal, Cell Signaling Tech. lot-4, 1:1000 dilution). Protein detection was carried out with horseradish peroxidase (HRP)-conjugated secondary antibodies, goat anti–mouse (Dako, P0447, 1:2500 dilution) or goat anti-rabbit IgG (Dako, P0448, 1:2500) and ECL substrate (GE Healthcare). Bands were quantified using Image J 1.32 software (v1.8, NIH). The intensity ratio of Parkin and actin, and Parkin and VDAC were calculated.

### Statistics

Data are represented as mean and standard error of the mean unless otherwise indicated in the Figure legends. Statistical significance was determined using ANOVA with Sidak’s or Bonferroni’s post-hoc multiple comparisons test and pair-wise comparisons were done with student paired t-test. Analysis was performed using Prism 7.0 software (GraphPad Inc.) and significance defined as p <0.05.

## ACKNOWLEDGMENTS

The authors’ work is supported by a Commonwealth studentship award funded by the UK government to MM (CSBD-2014-52., by the MRC SHIELD consortium (DHD PI)), MRNO2995X/1 and by an Innovation grant from the Antimicrobial Resistance Cross Council Initiative supported by the seven research councils (MR/M017931/1) to HMM. Electron microscopy support was provided through the University of Sheffield’s Faculty of Science Electron Microscopy Facility. PJS is supported as an NIHR Senior Investigator and by the NIHR Sheffield Biomedical Research Centre. Confocal microscopy support in Edinburgh was provided by the Queen’s Medical Research Institute Confocal and Advanced Light Microscopy Facility (QMRI CALM), and flow cytometry support in Edinburgh was provided by the QMRI Flow Cytometry and Cell Sorting Facility at the University of Edinburgh.

## AUTHOR CONTRIBUTIONS

MM, KBC, EF, ECP, CDR and JM conducted and analysed experiments. SPA and PJS provided expertise and access to equipment to facilitate Seahorse experiments. CP provided tissue culture and macrophage characterization expertise. KJdV provided expertise and methodology to enable imaging based analysis of fission. CJH and PB provided expertise in electron microscopy. AMC provided reagents and shared expertise in analysis of PI3K pathways. TJM provided expertise in microbiology and generated mutants used in experiments. HMM and DHD designed experiments and reviewed analysis with MM and KBC. MM, KBC, HMM, DHD wrote the paper with input from all authors.

## SUPPORTING INFORMATION

**Fig S1.**
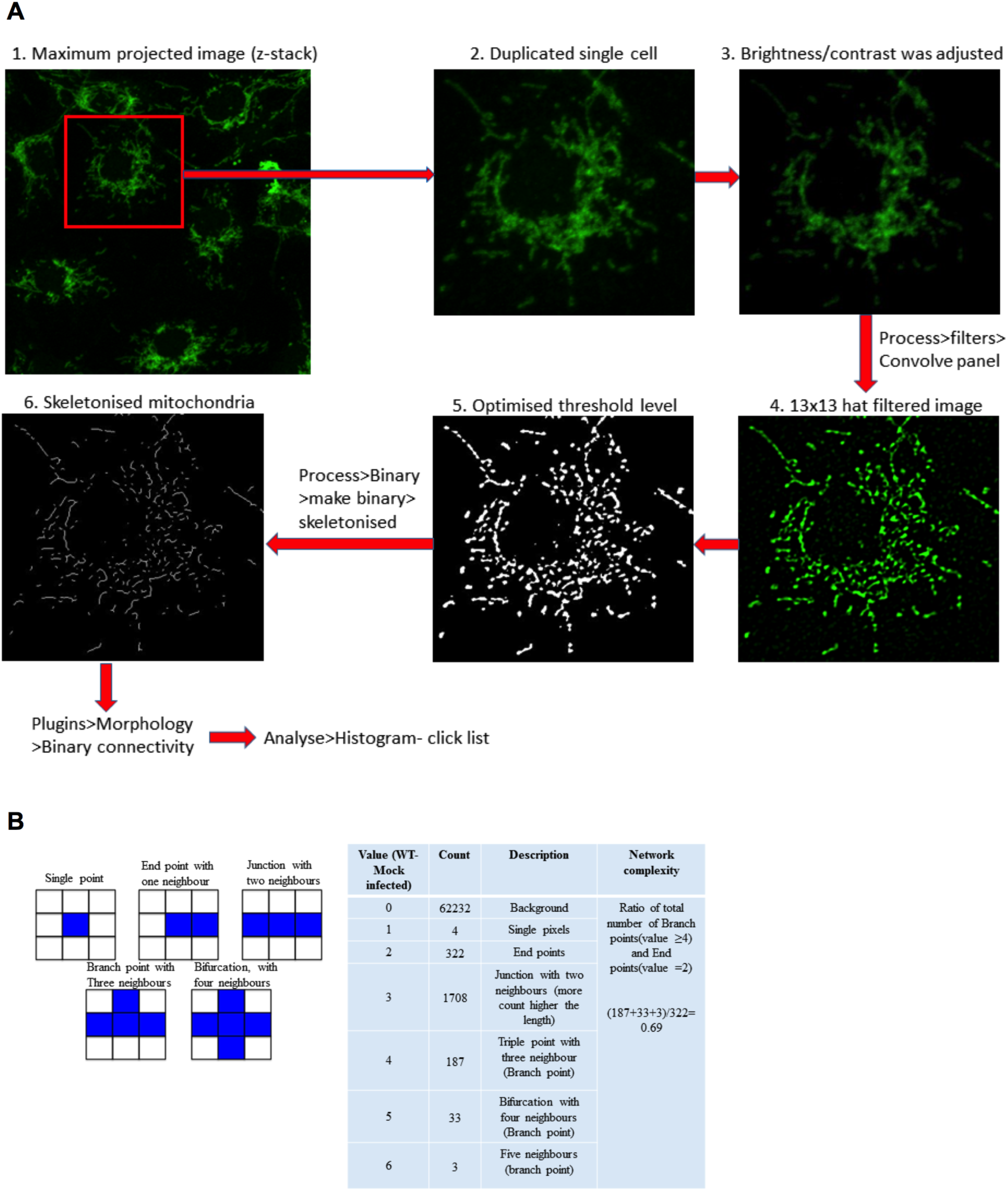
Schematic of calculation of mitochondrial network complexity using the binary connectivity and morphology plug-in of ImageJ. Schematic diagram illustrating method used to calculate the mitochondrial network complexity of the total population of mitochondria in each cell. Each cell was selected and duplicated from the maximum projected image. The resolution of each cell was enhanced using a 13 x 13 hat filter before threshold level correction was performed. The total population of mitochondria in each cell was skeletonized using a binary plugin. The voxel or mitochondrial connectivity was evaluated by the binary connectivity of morphology plug-in in ImageJ. The network complexity is defined as the ratio of total branch points and total end points. (B) Representative calculation of a single cell’s mitochondrial network complexity. The left rectangles show how the voxel connectivity were quantified using the binary connectivity plugin and the table defines each value used and how the ratio of total branch points and total end points was calculated from the binary pixels.

**Fig S2.**
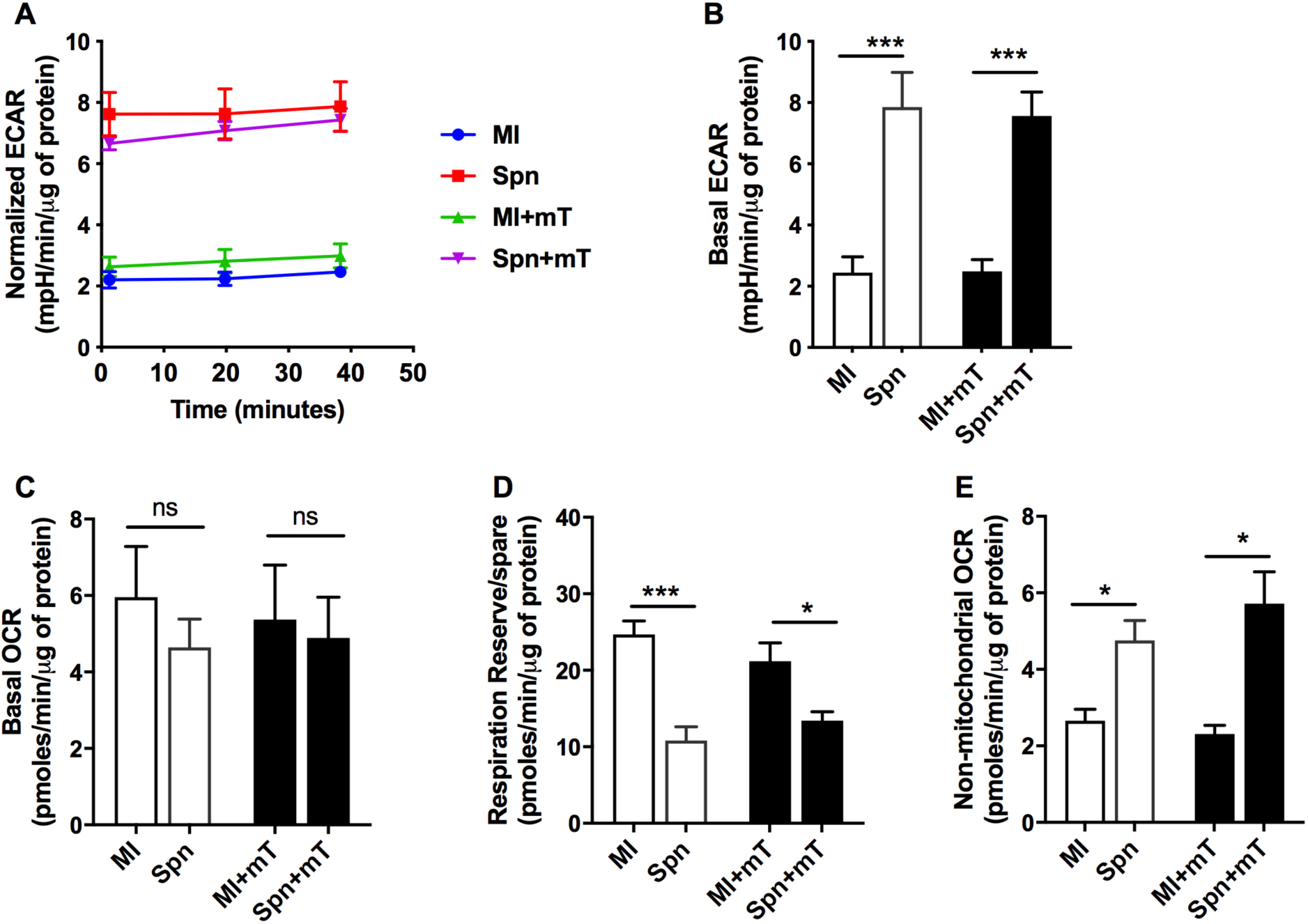
**Metabolic profile of BMDMs following *S. pneumoniae* challenge**. BMDMs were mock-infected (MI) or challenged with *S. pneumoniae* (Spn) for 12 h in the presence of pre-treatment with the mROS inhibitor MitoTempo (+mT) or vehicle control. Subsequently, the extracellular acidification rate (ECAR) and mitochondrial oxygen consumption rate (OCR) were measured by the Seahorse X24 extracellular flux analyser in the same experiments as Figure 3. The normalized ECAR (A), basal ECAR (B), basal OCR (C), respiration reserve (D) and non-mitochondrial OCR (E) were calculated. Data are shown as mean ± SEM, n=4. Statistical analysis was performed with one-way ANOVA with Sidak’s post-hoc test for multiple comparisons, *p≤0.05, ***p ≤0.001.

**Fig S3.**
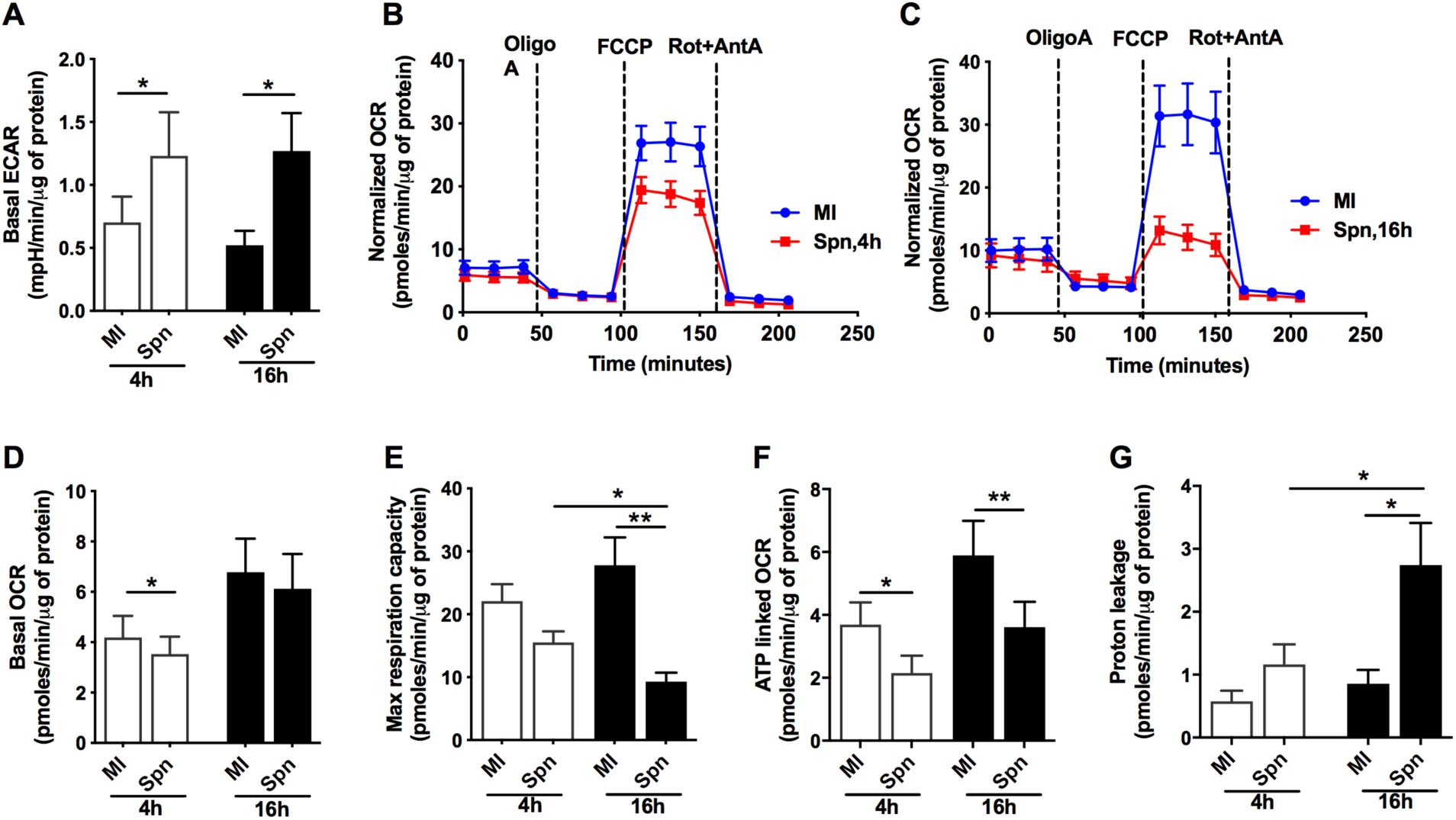
Metabolic profile of MDMs following *S. pneumoniae* challenge. Human monocyte-derived macrophages (MDMs) were mock-infected (MI) or challenged with *S. pneumoniae* (Spn) for 4 or 16 h and the extracellular acidification rate (ECAR) and mitochondrial oxygen consumption rate (OCR) was measured by the Seahorse X24 extracellular flux analyser. (A) Basal ECAR, (B) representative plots for the OCR kinetic data at 4 h and (C) at 16 h, (D) basal OCR, (E) maximum respiration capacity, (F) ATP-linked OCR and (G) proton leakage were calculated. Data are shown as mean ± SEM, n=5 (4 h) and n=6 (16 h). Statistical analysis was performed with one-way ANOVA with Sidak’s post-hoc test for multiple comparisons, *p≤0.05, ***p ≤0.001.

**Fig S4.**
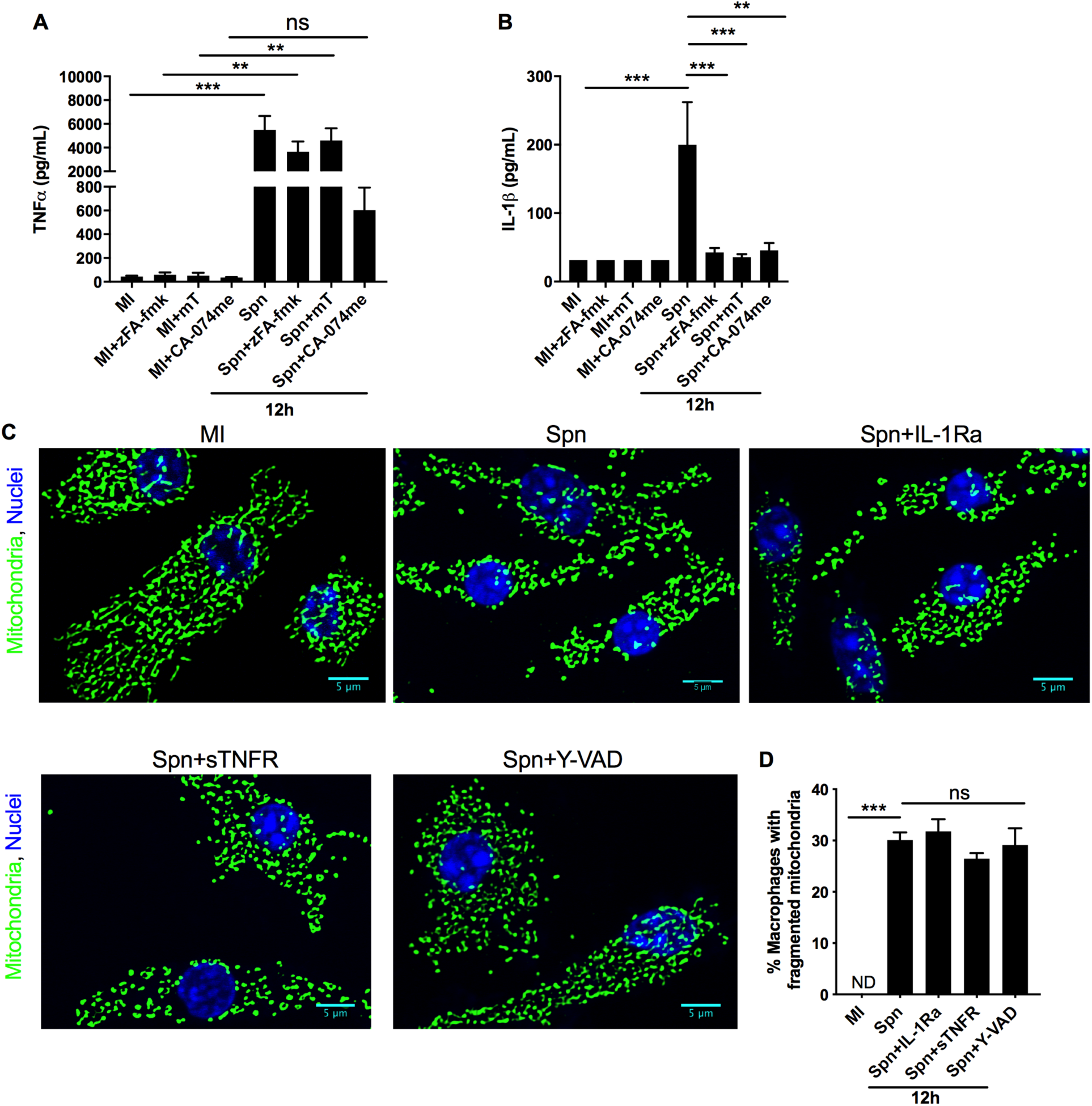
IL-1β production is regulated by mROS but does not regulate mitochondrial fission. BMDMs were mock-infected (MI) or challenged with *S. pneumoniae* (Spn) for 12 h in the presence or absence of pre-treatment with the cathepsin B inhibitors zFA-fmk or CA-047me or the mROS inhibitor MitoTempo (mT) and supernatants collected for assessment of (A) TNFα, n*=6* or (B) IL-1β by ELISA, n=. BMDMs were mock-infected (MI) or challenged with *S. pneumoniae* (Spn) for 12 h in the presence or absence of pre-treatment with IL-1RA, sTNFR1 or the caspase 1 inhibitor YVAD and cells stained with anti-TOMM20 and mitochondrial complexity calculated. (C) Representative filtered confocal images, from three independent experiments are shown and (D) the percentage of macrophages with fragmented mitochondria shown, n=3, scale bars = 5 µm. Data are shown as mean ± SEM. Statistical analysis was performed with one-way ANOVA with Sidak’s post-hoc test for multiple comparisons, *p≤0.05, **p ≤0.01. ***p ≤0.001.

**Fig S5.**
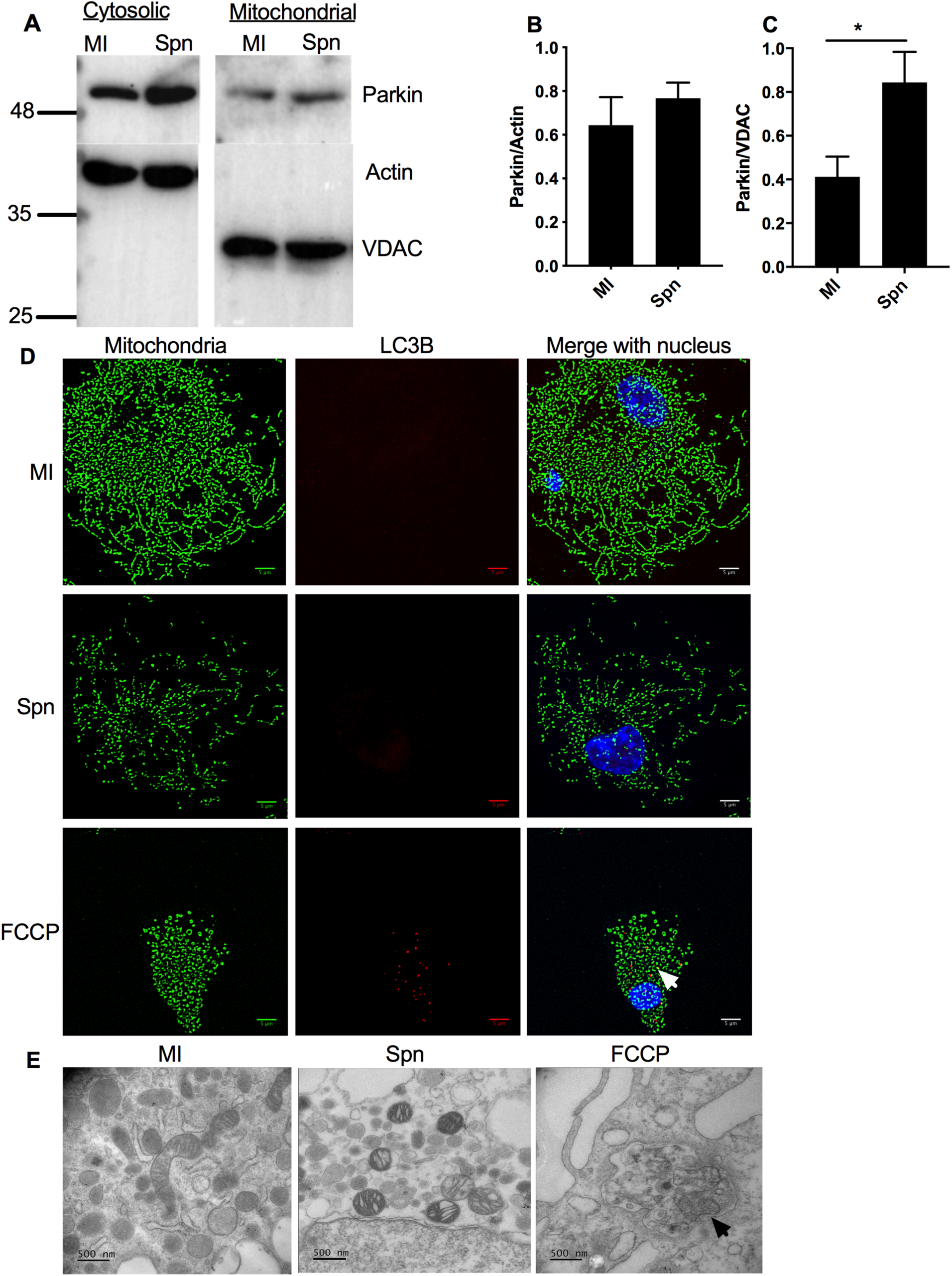
Parkin recruitment to mitochondria is not associated with mitophagy. BMDMs were mock-infected (MI) or challenged with *S. pneumoniae* (Spn) for 12 h and cells were lysed and fractionated into cytosolic and mitochondrial fractions. (A) A representative western blot after probing with anti-Parkin antibody is shown, with actin used as a cytosolic loading control and voltage dependent anion channel (VDAC) as a mitochondrial loading control. The blot is representative of three independent experiments. Densitometry was performed to estimate the Parkin/actin and (C) Parkin/VDAC ratios, n=3. Data are shown as mean ± SEM. Statistical analysis was performed with student paired *t* test for pair-wise comparisons. (D) BMDMs were MI or challenged with Spn or exposed to Carbonyl cyanide 4-(trifluoromethoxy) phenylhydrazone (FCCP; positive control). After 12 h BMDMs were harvested and stained with anti-TOMM20 (green) to outline mitochondrial structure or with anti-LC3B (red) as marker of mitophagy. A representative filtered image from three independent experiments is shown, scale bars = 5 µm. (E) Under the same conditions macrophages were harvested and examined by transmission electron microscopy to search for double membrane containing vacuoles containing mitochondria consistent with mitophagy (arrowhead), as shown with the positive control FCCP. The images are representative of three independent experiments. Scale bars =500 nm.

**Fig S6.**
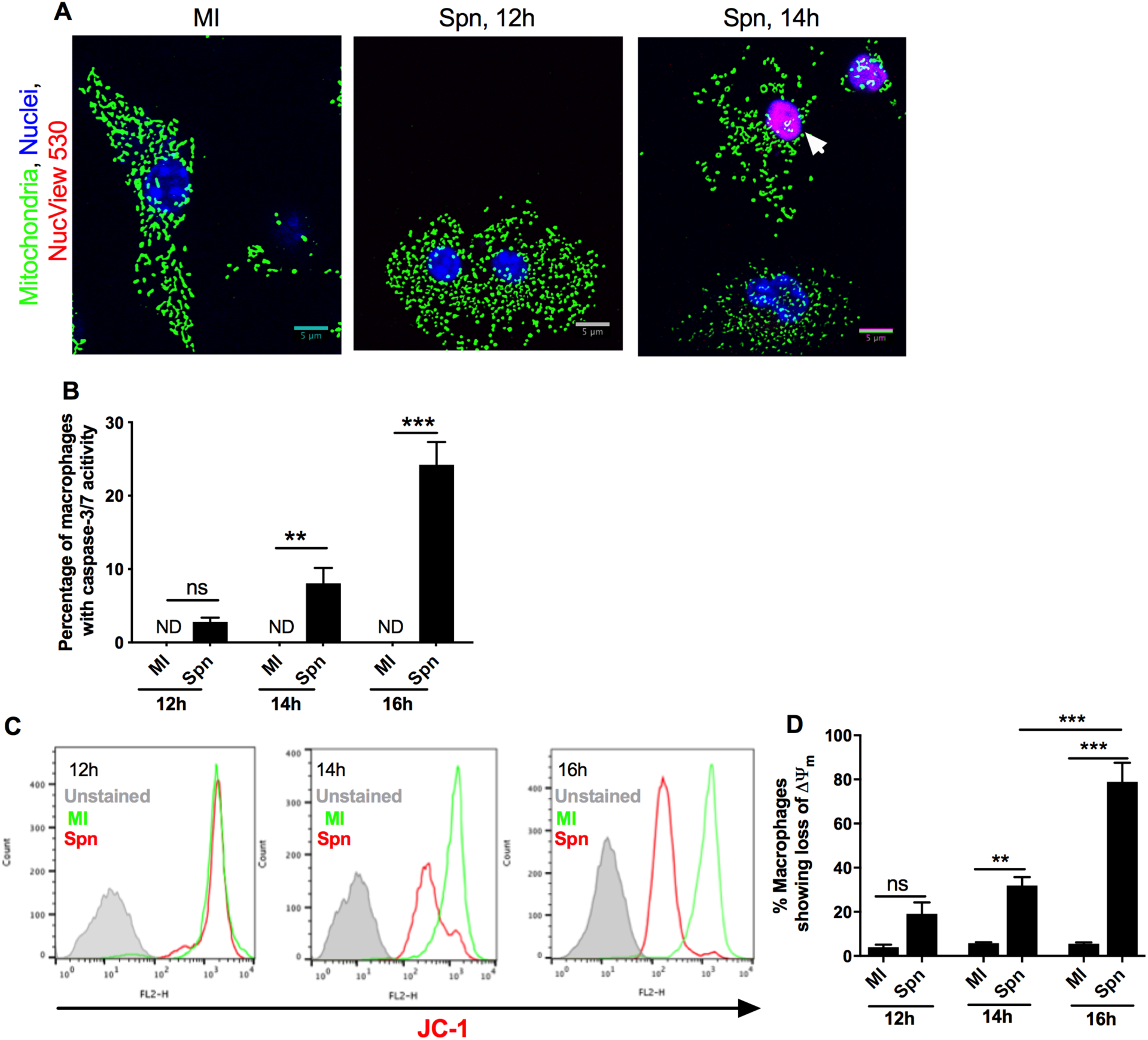
Apoptosis of macrophages following bacterial challenge occurs downstream of mitochondrial fission. BMDMs were mock-infected (MI) or challenged with *S. pneumoniae* (Spn) for 12-16 h and cells stained with NucView 530 (red) to detect caspase 3/7 activation during apoptosis and with DAPI (blue) to detect total cell count. Along with NucView red and DAPI staining, cells were also stained with anti-TOMM20 (green) (A) Representative filtered images from three independent experiments are shown, scale bars = 5 µm and (B) quantification of the percentage of caspase 3/7 positive BMDMs, n=3 are shown. Under the same conditions JC-1 staining was performed to measure loss of Δψ_m_ by flow cytometry. (C) Representative histograms, from three independent experiments, are shown at 12-16 h after challenge with unstained cells in grey, MI in green and Spn exposed BMDMs in red. (D) The percentage of BMDMs showing loss of Δψ_m_ at each time point are shown, n=3. Data are shown as mean ± SEM. Statistical analysis was performed with one-way ANOVA with Sidak’s post-hoc test for multiple comparisons, *p≤0.05, **p ≤0.01. ***p ≤0.001.

**Fig S7.**
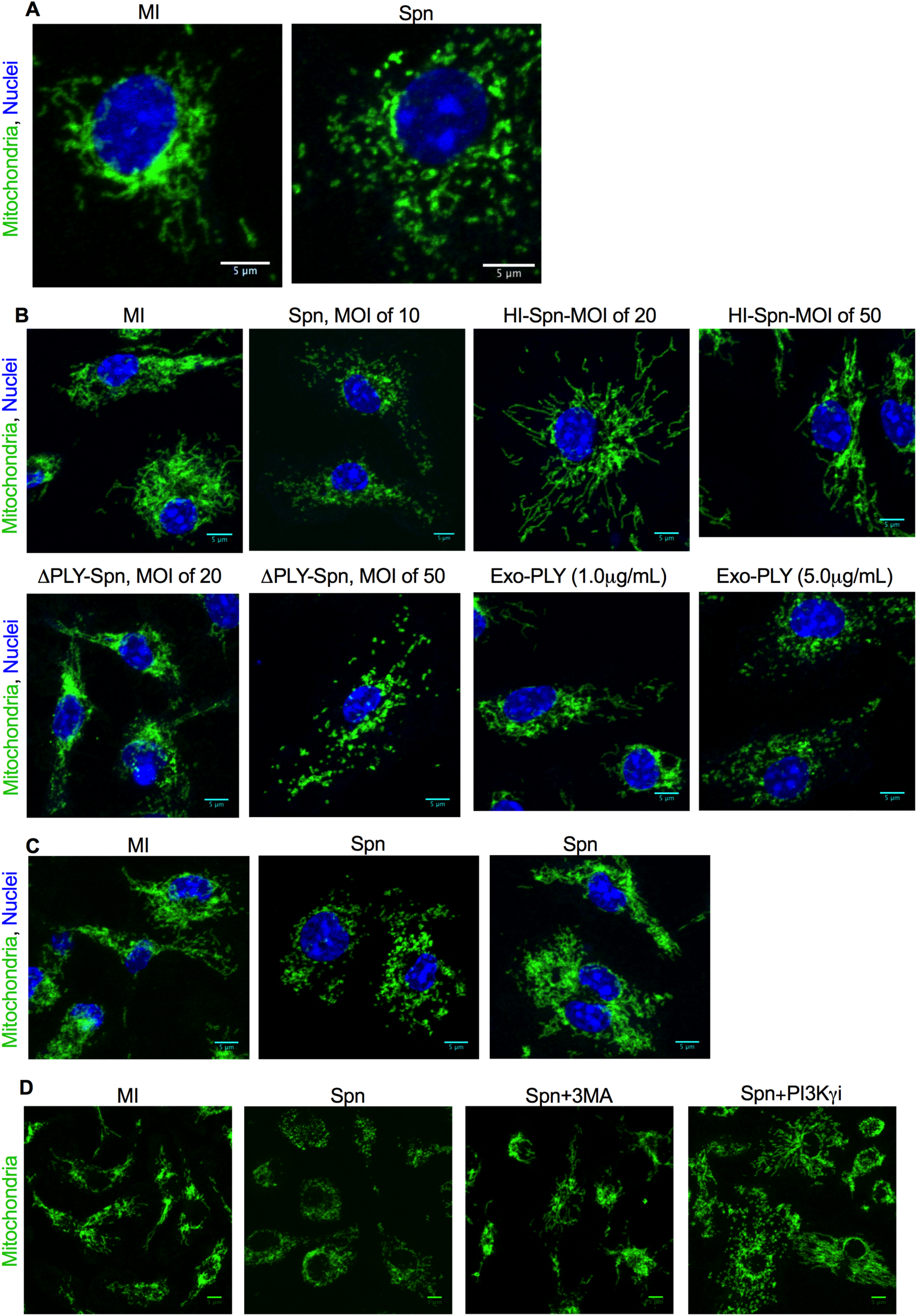
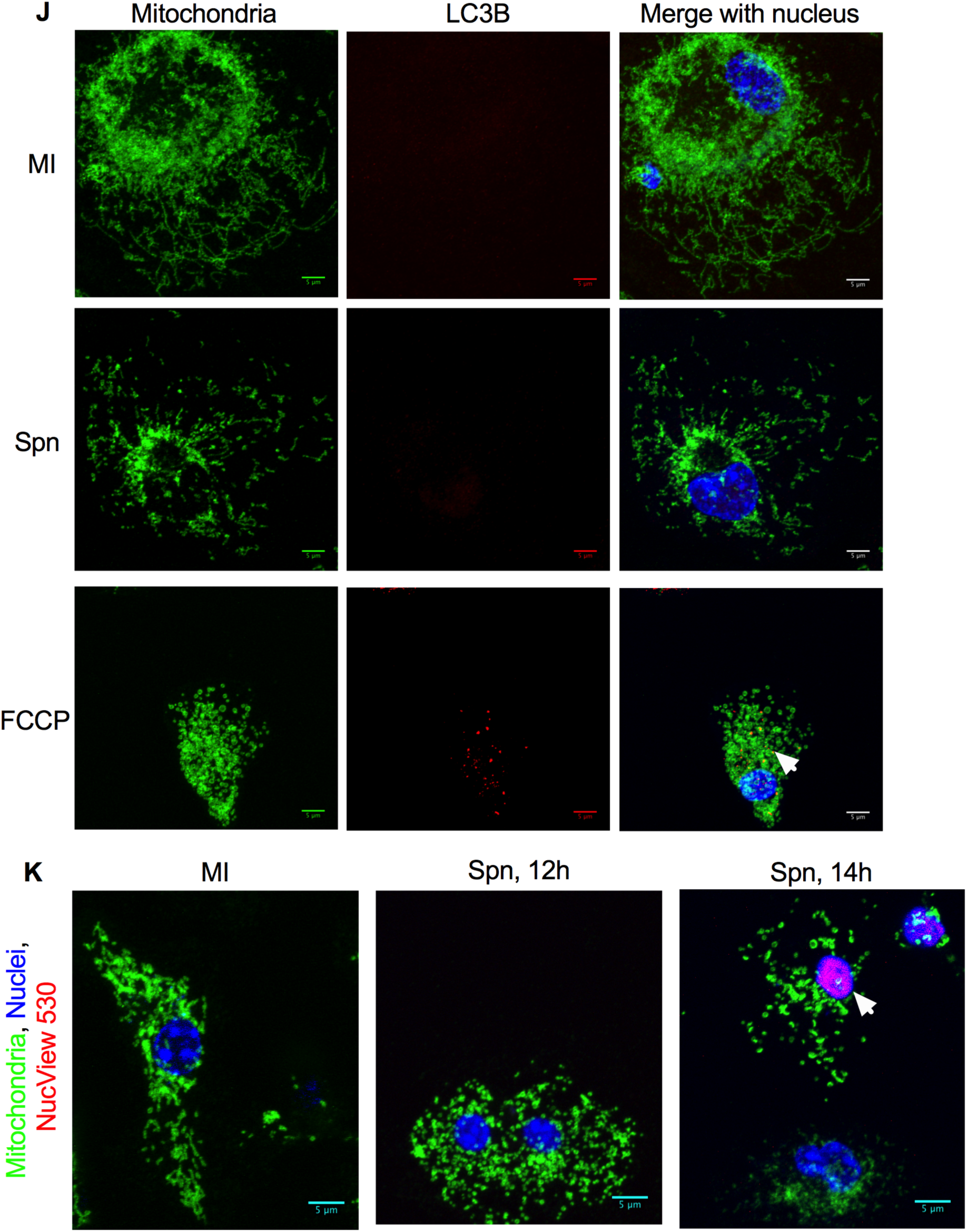
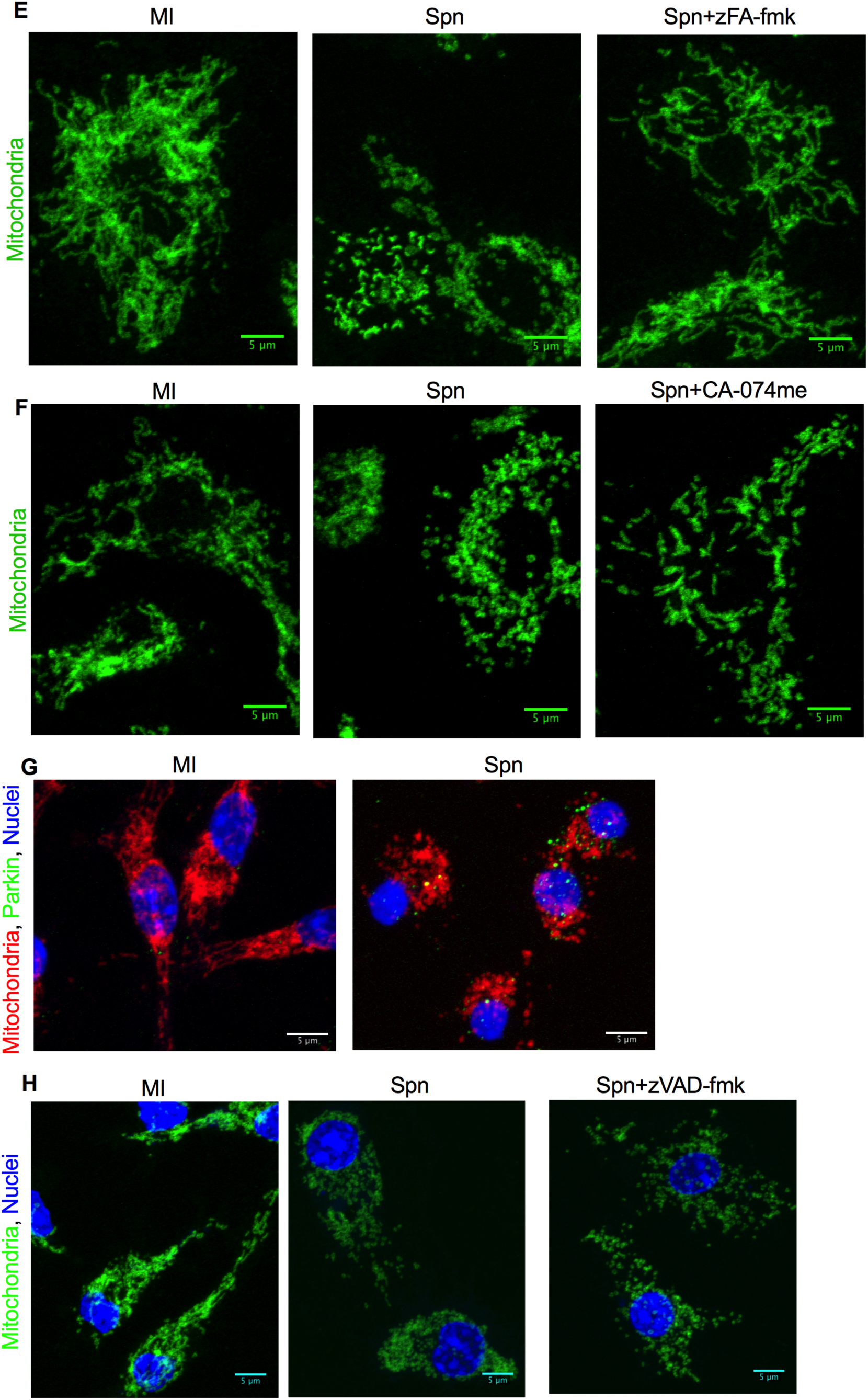
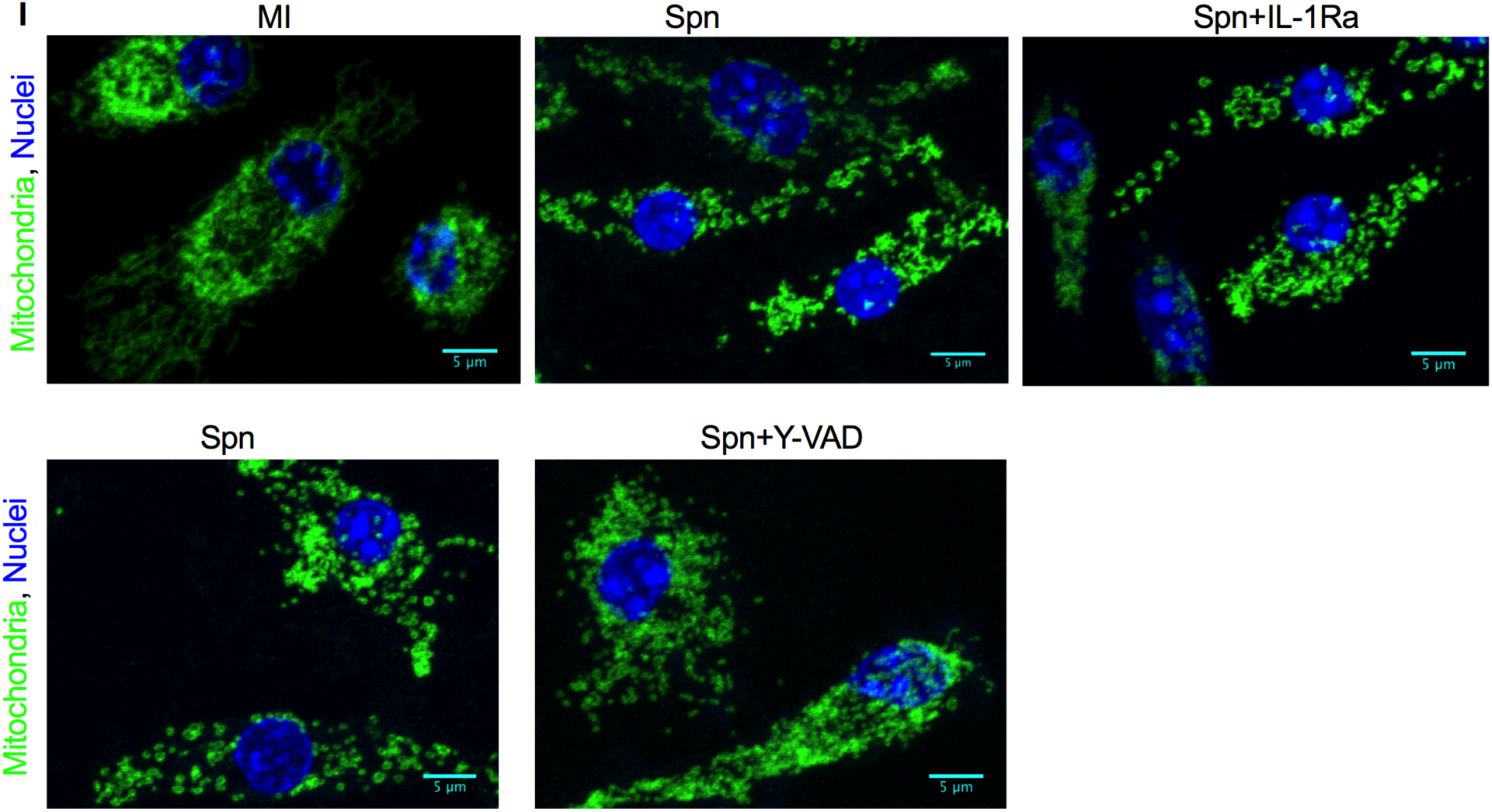
Unfiltered confocal images used to generate the filtered images shown in the manuscript. Images were generated using bone marrow derived macrophages (BMDMs) that were mock-infected (MI) or exposed to *S. pneumoniae* (Spn) at a MOI of 10, for 12 h, unless otherwise stated, in the presence or absence of various inhibitors and vehicle controls and then stained as described in each individual Figure legend. Representative unprocessed images generated by confocal microscopy are depicted and represent the images used to generate the indicated filtered images shown throughout the text. Scale bars = 5 µm unless otherwise stated (A) MI or Spn BMDM stained with the mitochondrial outer membrane specific marker TOMM20 (green) to delineate mitochondrial structure. The unprocessed images correspond to the images Fig. 1A. (B) MI or Spn BMDMs or those exposed to heat-inactivated Spn (HI-Spn) or a pneumolysin deficient Spn mutant (ΔPLY Spn), at the indicated multiplicity of infection, or exogenous pneumolysin (0.5 µg/mL-5 µg/mL) were stained with anti-TOMM20 (green) or Draq5 staining to label nuclei (blue). Unprocessed images represent the images as shown in Fig. 2A. (C) MI or Spn BMDM were pre-treated with mROS inhibitor MitoTempo (+mT) or vehicle control before staining with anti-TOMM20 (green) and Draq5 (blue). Unprocessed images represent the images as shown in Fig. 3H. (D) MI or Spn BMDMs were challenged in the presence of vehicle control or the indicated PI3K inhibitors (3 Methyladenine (3MA), Ly294002 (Pan-PI3Ki), and selective inhibitors of PI3Kα (PI3Kαi), PI3Kγ (PI3Kγi) or PI3Kδ (PI3Kδi) isoforms and stained with anti-TOMM20 (green) staining. Representative unprocessed images represent the images shown in Fig. 4D. (E,F) MI or Spn BMDMs were challenged in the presence of vehicle control, zFA-fmk (E) or CA-074me (F). Cells were stained with anti-TOMM20 (green) and representative unprocessed images represent the images shown in Fig. 5E (E) and Fig. 5F (F). (G) MI or Spn BMDMs were stained with anti-TOMM20 (red), Draq5 (blue) and anti-Parkin antibody (green). Representative unprocessed confocal images correspond to Fig. 6A. (H) MI or Spn BMDMs in the presence or absence of zVAD-fmk were stained with anti-TOMM20 (green). Representative unprocessed confocal images correspond to Fig. 6D. (I) MI or Spn BMDMs in the presence or absence of pre-treatment with IL-1RA, sTNFR1 or the caspase 1 inhibitor YVAD were stained with anti-TOMM20 (green) and Draq5 (blue). Representative unprocessed images were those used in Fig. S5C. (J) MI or Spn BMDMs or BMDM exposed to FCCP (positive control) were stained with anti-TOMM20 (green), Draq5 (blue) or with anti-LC3B (red). Representative unprocessed images correspond to those used in Fig. S6D. (K) MI or Spn BMDMs at the indicated time points were stained with anti-TOMM20 (green), NucView 530 (red) to detect caspase 3/7 activation during apoptosis and with DAPI (blue) to stain nuclei. Representative unprocessed images correspond to those used in Fig. S7A.

